# mitoNEET Regulates Mitochondrial Iron Homeostasis Interacting with Transferrin Receptor

**DOI:** 10.1101/330084

**Authors:** Takaaki Furihata, Shingo Takada, Satoshi Maekawa, Wataru Mizushima, Masashi Watanabe, Hidehisa Takahashi, Arata Fukushima, Masaya Tsuda, Junichi Matsumoto, Naoya Kakutani, Takashi Yokota, Shouji Matsushima, Yutaro Otsuka, Masaki Matsumoto, Keiichi I. Nakayama, Junko Nio-Kobayashi, Toshihoko Iwanaga, Hisataka Sabe, Shigetsugu Hatakeyama, Hiroyuki Tsutsui, Shintaro Kinugawa

## Abstract

Iron is an essential trace element for regulation of redox and mitochondrial function, and then mitochondrial iron content is tightly regulated in mammals. We focused on a novel protein localized at the outer mitochondrial membrane. Immunoelectron microscopy revealed transferrin receptor (TfR) displayed an intimate relationship with the mitochondria, and mass spectrometry analysis also revealed mitoNEET interacted with TfR *in vitro*. Moreover, mitoNEET was endogenously coprecipitated with TfR in the heart, which indicates that mitoNEET also interacts with TfR *in vivo*. We generated mice with cardiac-specific deletion of mitoNEET (mitoNEET-knockout). Iron contents in isolated mitochondria were significantly increased in mitoNEET-knockout mice compared to control mice. Mitochondrial reactive oxygen species (ROS) were higher, and mitochondrial maximal capacity and reserve capacity were significantly decreased in mitoNEET-knockout mice, which was consistent with cardiac dysfunction evaluated by echocardiography. The complex formation of mitoNEET with TfR may regulate mitochondrial iron contents via an influx of iron. A disruption of mitoNEET could thus be involved in mitochondrial ROS production by iron overload in the heart.

## Introduction

Mitochondrial function is impaired with aging in various organs, including the heart, brain, and skeletal muscle (Bagh, Thakurta et al., 2011, Kwong & Sohal, 2000, Li, Kumar Sharma et al., 2013, Sugiyama, Takasawa et al., 1993). Age-dependent mitochondrial dysfunction is one of the main causes of organ failure and cellular dysfunction (Dai & Rabinovitch, 2009). Mitochondria are one of the main sources of reactive oxygen species (ROS) generation, which is associated with mitochondrial dysfunction. Conversely, mitochondria are highly susceptible to attack by ROS, because they contain iron/sulfur clusters (ISCs). Thus, once the mitochondria are impaired for any reason, an increase in ROS generation can, in itself, induce a vicious cycle of further impairment in mitochondria function, further ROS generation, and cellular injury (Ikeuchi, Matsusaka et al., 2005, Suematsu, Tsutsui et al., 2003).

Iron is one of the most abundant metals on earth and an essential trace element for both cellular energy and metabolism and the maintenance of body homoeostasis. Excessive iron damages cells via iron toxicity and ROS production, while iron deficiency impairs cellular proliferation (Oliveira, Rocha et al., 2014). Thus iron concentrations in mammals need to be tightly and constantly regulated at the dietary, plasma, extracellular, cellular and mitochondrial levels (Ganz, 2013). Despite the numerous studies on iron homeostasis in the last few decade (Anderson, Shen et al., 2012, Anderson & Vulpe, 2009), mitochondrial iron homeostasis remains largely unexplored. ISCs and heme are produced only within mitochondria (Richardson, Lane et al., 2010), and are used as components of many enzymes in the cytosol and mitochondria. Therefore, iron-homeostasis regulation is closely associated with mitochondrial function and cellular function. Some proteins involved in influx of iron to mitochondria, transfer of iron within mitochondria, and efflux of iron from mitochondria have been identified, and all these proteins exist in the mitochondrial inner membrane or within mitochondria (Branda, Cavadini et al., 1999, Ichikawa, Bayeva et al., 2012, Paradkar, Zumbrennen et al., 2009, Srinivasan, Pierik et al., 2014, Vigani, Tarantino et al., 2013). Iron bound on transferrin (Tf) and transferrin receptor (TfR), namely Tf-TfR complex, is carried to the cytosol as endosomes, and then iron dissociated from the complex transports across the endosomal membrane to the labile iron pool in the cytosol. Iron from the labile iron pool is carried into the mitochondria to utilize this iron for mitochondrial iron homeostasis (Gkouvatsos, Papanikolaou et al., 2012). Whereas some proteins involved in iron inflow into the mitochondria are known to be in the inner mitochondrial membrane (Richardson et al., 2010), novel proteins involved in mitochondrial iron homeostasis may also exist in the outer mitochondrial membrane (OMM).

It has recently been reported that mitoNEET is a novel protein localized at the OMM and a target protein of the insulin-sensitizing drug pioglitazone (Wiley, Murphy et al., 2007a). Analysis of the expression of mRNAs from murine tissues revealed that mitoNEET was widely expressed in mice, with especially high levels in the heart.

Experiments using optical and electron paramagnetic resonance spectroscopy clarified that mitoNEET contained iron/sulfur clusters that were redox-active and functioned as electron-transfer proteins (Wiley, Paddock et al., 2007b). Moreover, complex I-linked state 3 respiration was significantly decreased in isolated mitochondria from the hearts of mitoNEET-null mice. These results allowed us to hypothesize that mitoNEET was a candidate for the regulatory protein of iron homeostasis.

The regulation of mitochondrial function is especially important for the maintenance of cardiac function, because the heart is a mitochondria-rich organ (Rosca & Hoppel, 2010). In the present study, we focused on the mechanisms underlying the regulation of mitochondrial iron homeostasis in the heart, and verified our hypothesis that mitoNEET is the regulator of iron homeostasis. To accomplish this, we used lox-P and homologous recombination to establish mice with cardiac-specific deletion of mitoNEET.

## Results

### mitoNEET combines with TfR and is regulated by iron

TfR have been known to be located in plasma membrane or endosome (Das, Nag et al., 2016). Surprisingly, in mitochondrial fraction, TfR were detected evaluated by immunoblot, though there existed no other proteins localized in cellular matrix (glyceraldehyde phosphate dehydrogenase; GAPDH), endosome (Ras-related protein Rab-5; Rab5) or plasma membrane (Cadherin) (**Fig 1A**). The silver-intensified immunogold method for electron microscopy revealed the subcellular localization of TfR in the C2C12 cells incubated with iron. Immunogold particles showing the localization of TfR were frequently associated with (cytoplasmic) vesicles and vacuoles of various sizes and shapes, which might correspond to endosomes. In addition, gold particles displayed an intimate relationship with mitochondria, where they attached the outer membrane of mitochondria. (**Fig 1B**). Protein-protein interactions are often important to realize biological functions *in vivo*. To elucidate what kind of mitochondrial protein is involved in mitochondrial iron homeostasis, we focused on TfR. That is because TfR is thought to have a critical role in iron uptake into the cell. So silver-stained gels with the immunoprecipitated protein by TfR antibody in the mitochondria from the mouse heart acquired some bands (**Fig 1C**), suggesting that endogenous TfR coprecipitated with some endogenous mitochondrial proteins. We think that mitoNEET could play a critical role in mitochondrial iron homeostasis, for mitoNEET simultaneously fulfill below conditions; localized at the OMM (Das et al., 2016, Wiley et al., 2007a), containing ISCs (Paddock, Wiley et al., 2007), and involved in iron metabolism (Kusminski, Holland et al., 2012) (**Fig 1D**). Actually, candidate proteins were picked up by using mass spectrometry, and what we focused on as the strongest candidate was just TfR (**Fig 1E, Appendix Table S1, S2**). Moreover, mitoNEET was endogenously coprecipitated with TfR in the heart, which indicates that mitoNEET also interacts with TfR *in vivo* (**Fig 1F, 1G**). The levels of mitoNEET protein expression were higher in the addition of ferric ammonium citrate, and lower in the addition of desferioxamine compared to control (**Fig 1H**), and the changes in iron content were consistent with the changes in mitoNEET protein expression (**Appendix Fig S1**), suggesting that there may be some connections between mitoNEET and iron homeostasis. There was a significant positive correlation between mitoNEET expression levels and mitochondrial respiration (**Fig 1I**), suggesting that mitoNEET may have an important role in regulating cellular function via iron homeostasis.

**Figure 1.**
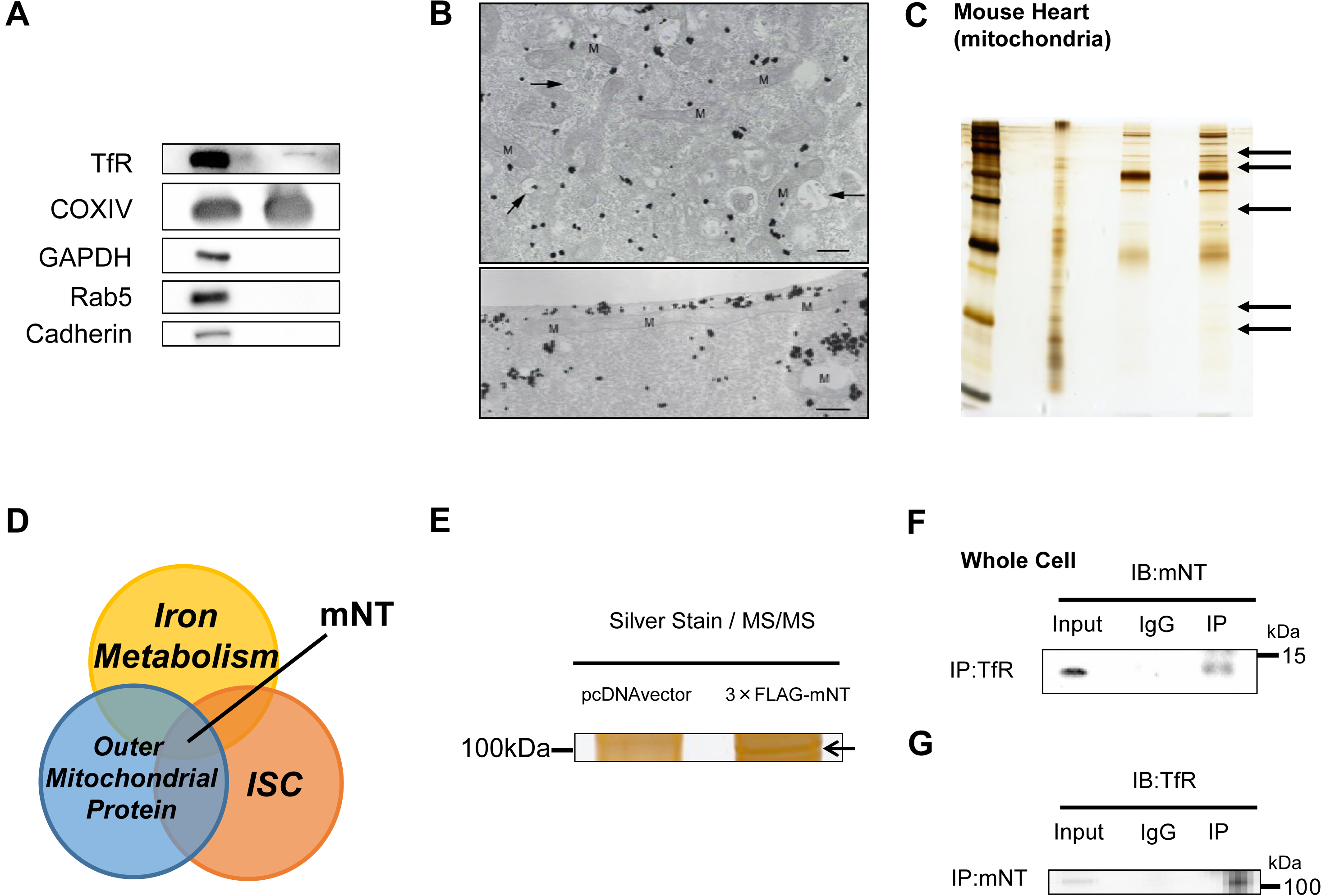
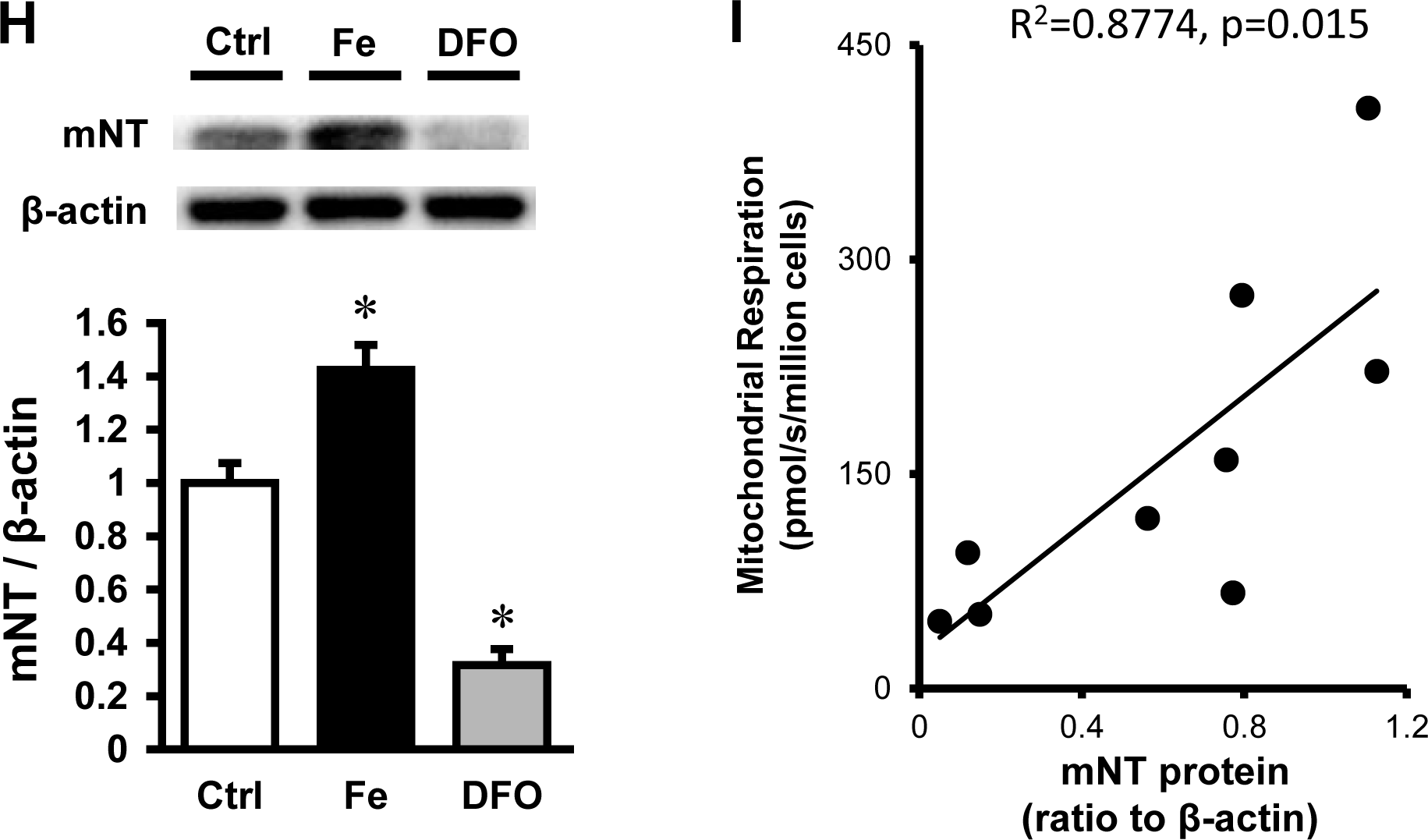
The Role of mitoNEET in Mitochondrial Iron Homeostasis. (A) Immunoblot of supernatant and pellet from mitochondrial fractions of the mouse heart. (B) Electron microscopy of TfR in C2C12 cells. In upper panel, Gold particles, black dots in figures, showing the existence of TfR are broadly distributed throughout the cytoplasm but tend to be condensed around vacuoles (arrows) and mitochondria. Lower panel shows an intimate relationship of gold particles with a long mitochondrion. Bar, 0.5 µm. (C) Silver-stained gels with the immunoprecipitated protein by TfR antibody in the mitochondrial protein from the mouse heart (black arrows). (D) Venn diagram showing the central role of mitoNEET in new mitochondrial iron regulation involved in iron metabolism, outer mitochondrial proteins, and iron sulfur-cluster(ISC). Silver-stained gels with the immunoprecipitated protein from HEK 293 cells transfected with and without 3×;FLAG-mitoNEET containing the pcDNA3 promoter. Interaction of endogenous TfR and mitoNEET. The whole cell lysate of the mouse heart was subjected to immunoprecipitation with anti-TfR antibody or normal rabbit IgG followed by immunoblot with mitoNEET antibody. An input representing 5 µg of the whole cell lysate was used for each immunoprepicitation. (G) Interaction of endogenous mitoNEET and TfR. (H) Representative immunoblot and summary data of mitoNEET protein expression normalized to beta-actin in mouse C2C12 cells in the addition of iron or the reduction of iron with DFO compared to control. Data are shown as the mean ± SE. n=7. *P<0.05 vs. Ctrl. (I) Correlation between mitochondrial respiration and mitoNEET protein expression normalized to beta-action. WCL, whole cell lysate; IB, immunoblot; IP, immunoprecipitation; TfR, transferrin receptor; mNT, mitoNEET; M, mitochondria; Ctrl, control; Fe, the addition of ferric ammonium citrate; DFO, desferioxamine.

### mitoNEET and Tranferrin Receptor Colocalizes on the Mitochondria

If mitoNEET interact with transferrin receptor, these proteins should colocalize in the mitochondria. We subsequently performed immunoprecipitation by using mitochondrial fraction. As we expected, mitoNEET coprecipitated with TfR in mitochondria from the heart in mice, suggesting that TfR, which has been thought to be only plasma membrane or endosomes, did exist with mitochondria (**Fig 2A**). Moreover, to determine whether TfR localizes to the OMM, we used digitonin, which is efficiently able to extract cholesterol in the OMM protein (Arasaki, Shimizu et al., 2015). To confirm the purity of isolated mitochondria, Cadherin and GAPDH were analyzed by immunoblot. These were not detected in supernatant and pellet. Heat-shock protein 60 was not detected in supernatant even with the treatment of digitonin, suggesting that digitonin used in the present study did not affect the inner mitochondrial membrane. Voltage-dependent anion channel (VDAC), a known OMM protein, was dose-dependently increased in supernatant and decreased in pellet by digitonin treatment. The changes in mitoNEET and TfR protein levels by the treatment with digitonin were similar to VDAC (**Fig 2B**). These data strongly suggest that TfR localizes to the OMM interacting with mitoNEET, besides plasma membrane or endosome. Immunostaining for mitoNEET (green) merged with TfR (red) in mouse C2C12 cells. In control, mitoNEET was localized at the OMM along surface of the mitochondria, and TfR mainly existed on the inside of cytosolic membrane. In iron overload circumstances, TfR partially transferred from membrane to cytosol, some of which colocalized with mitoNEET, suggesting that this increase of colocalization helps TfR to stay on the mitochondria in order to cope with iron overload (**Fig 2C**).

**Figure 2.**
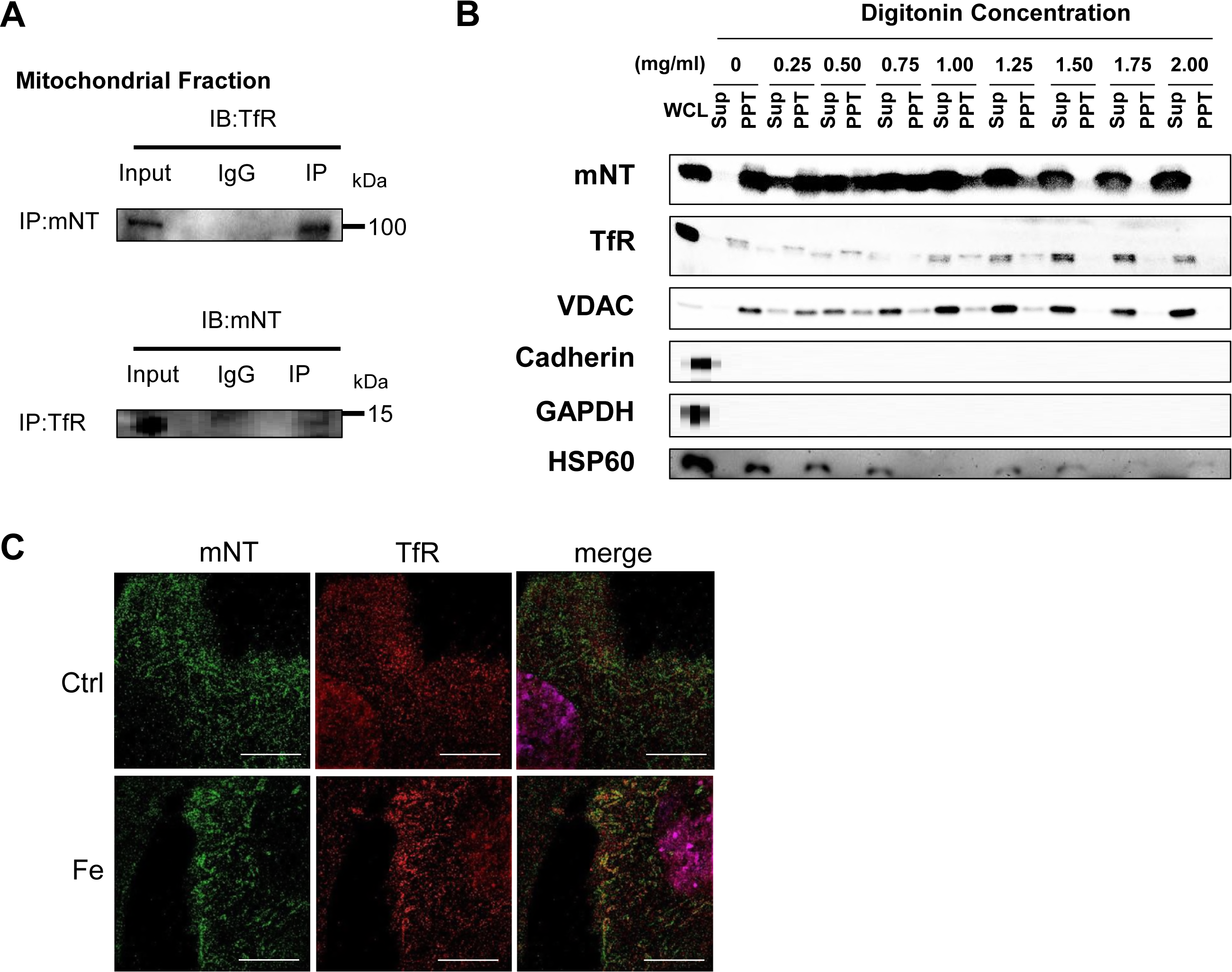
The Colocalization of mitoNEET and Transferrin Receptor on the Outer Mitochondrial Membrane. (A) Interaction of mitoNEET and TfR with mitochondria of the mouse heart. An input representing 100 µg of the mitochondria lysate was used. (B) Immunoblot of supernatant and pellet from mitochondrial fractions after digitonin treatment with the indicated concentrations. (C) Representative images obtained from structured illumination microscopy of C2C12 cell stained for mitoNEET and TfR. mitoNEET (green) and TfR (red) visualized using Alexa Fluoro Plus 488 and Alexa Fluoro Plus 555. Left, C2C12 cell imaged with 488 nm filter. Center, C2C12 cell imaged with 555 nm filter. Right, merged image. Bar, 10 μm. IB, immunoblot; IP, immunoprecipitation; KO, knockout; mNT, mitoNEET; TfR, transferrin receptor; ANT, adenine nucleotide translocator; WCL, whole cell lysate; Sup, supernatant; PPT, pellet; VDAC, voltage-dependent anion channel; HSP60, heat-shock protein 60; Ctrl, control; Fe, the addition of ferric ammonium citrate.

### Creation of Cardiac-Specific mitoNEET-knockout Mice

To establish mice with cardiac-specific deletion of mitoNEET, lox-P and homologous recombination strategies were used **(Appendix Fig S2A)**. As controls, we generated mitoNEET flox/flox mice in similar manner. Cardiac-specific deletion of mitoNEET was achieved using αMHC-Cre (**Appendix Fig S2B**). The cardiac-specific mitoNEET-knockout mice were viable and fertile, and there were no differences in appearance, body weight or cardiac phenotype between control and mitoNEET-knockout mice at a young age (about 3 months). Levels of the mRNA of *CISD1*, encoding the mitoNEET protein, were significantly lower in the whole heart of mitoNEET-knockout than control mice (**Appendix Fig S2C**). To characterize the mitoNEET protein expression, we generated a mitoNEET polyclonal antibody using a C-terminal fragment of mitoNEET as an antigen. As shown in **Appendix Fig S2D**, immunoblot, analyzed by Tris-Tricine sodium dodecyl sulfate polyacrylamide gel electrophoresis (SDS-PAGE), confirmed the mitoNEET expression in the hearts of control mice. The C-terminal fragment of mitoNEET (below 2 kDa) is also shown as a positive control. As expected, immunoblot analysis revealed that mitoNEET protein expression was widely detected in the brain, heart, liver, kidney and skeletal muscle of control mice, but was absent in the hearts of mitoNEET-knockout mice (**Fig 3A**). Immunohistochemical analysis of mitoNEET expression indicated that mitoNEET was broadly present in cardiac cells (**Appendix Fig S2E**).

**Figure 3.**
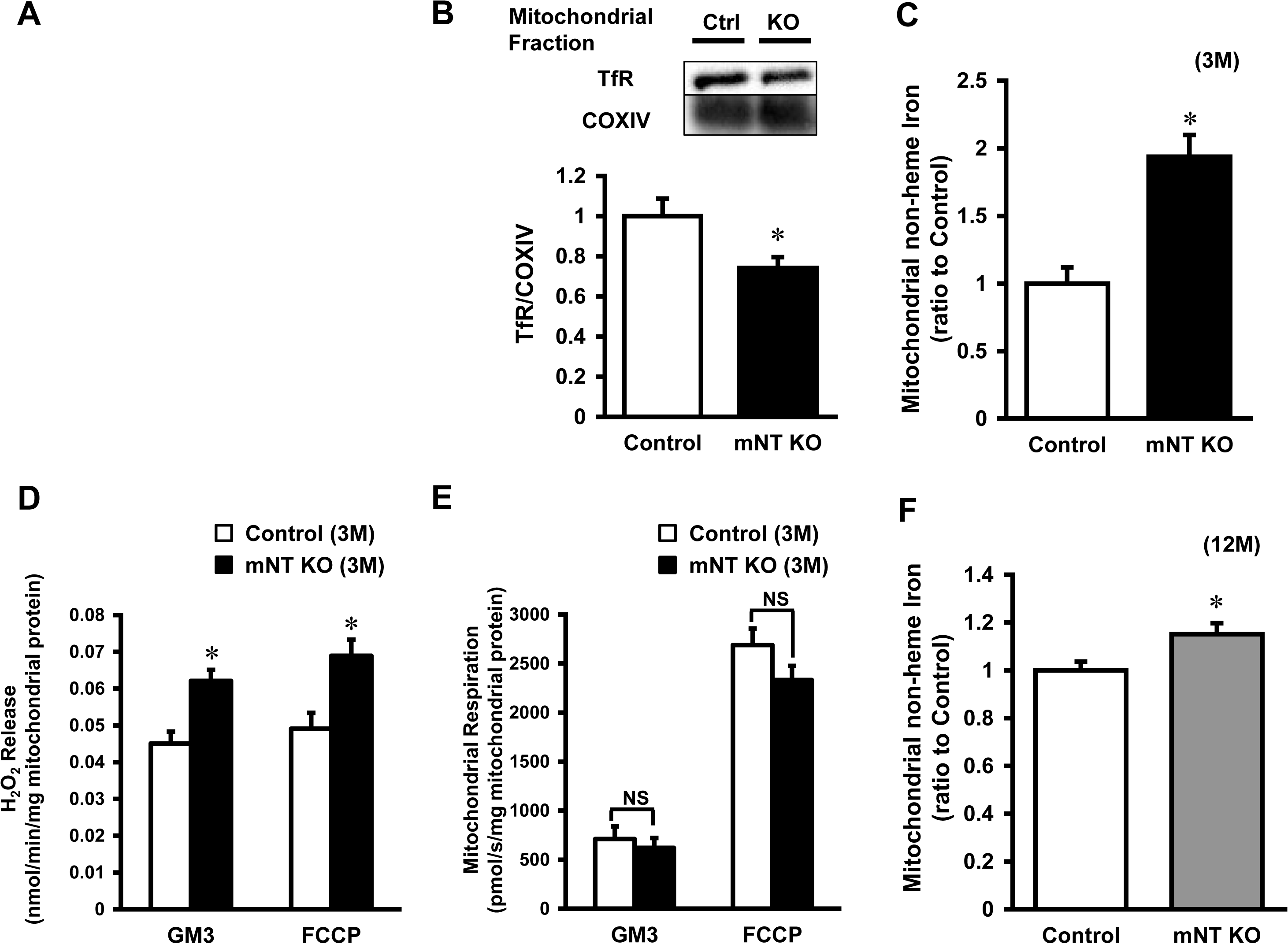
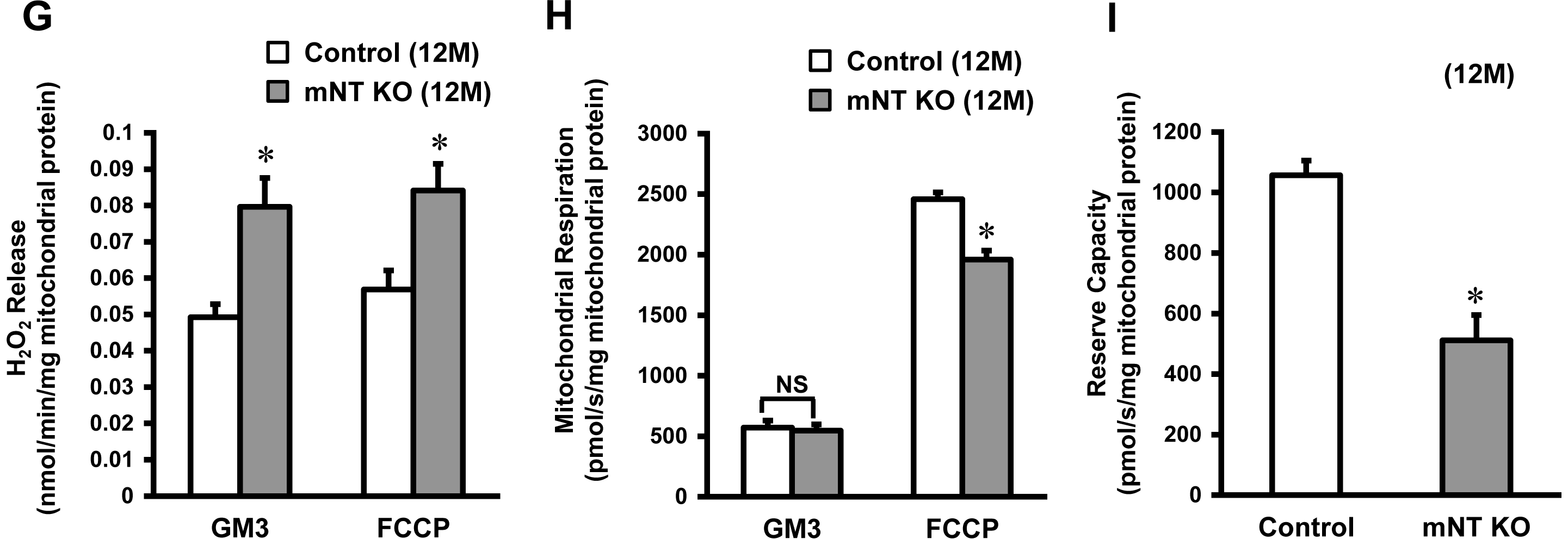
Mitochondrial Iron, ROS, and Respiration in the Heart of mitoNEET-knockout mice. (A) Representative immunoblot of the mitoNEET protein in various organs from mitoNEET-knockout mice and control mice. (B) Representative immunoblot of the transferrin receptor protein from control mice (n=6) and mitoNEET-konckout (n=6) mice in the mitochondrial fraction of the heart normalized to COX IV. (C) Levels of mitochondrial iron contents in 3-month-old mitoNEET-knockout mice relative to control mice (n=8-9). (D) H_2_O_2_ release originating from isolated mitochondria in the heart of 3-month-old control mice (n=5) and 3-month-old mitoNEET-knockout mice (n=5) during state 3 respiration with complex I-linked substrates (left bars) and in maximal capacity of an electron-transfer system with FCCP (right bars). (E) Mitochondrial respiration in isolated mitochondria from the heart of 3-month-old mice during state 3 with complex I-linked substrates (left bars) and maximal capacity of the electron-transfer system with FCCP (right bars). (F) Levels of mitochondrial iron contents in 12-month-old mitoNEET-knockout mice relative to 12-month-old control mice (n=11-14). (G) H_2_O_2_ release in 12-month-old control mice (n=5) and 12-month-old mitoNEET-knockout mice (n=5). (H) Mitochondrial respiration in isolated mitochondria from the heart of 12-month-old mice. (I) Reserve capacity in 12-month-old control mice (left bar) and mitoNEET-knockout mice (right bar). mNT, mitoNEET; GAPDH, glyceraldehyde phosphate dehydrogenase; B, brain; H, heart; Li, liver; K, kidney; SM, skeletal muscle; COX, cytochrome c oxidase; M, month. GM3, state 3 respiration with glutamate and malate; FCCP, carbonyl cyanide-p-trifluoromethoxyphenylhydrazone.

### Interaction between mitoNEET and TfR in mitoNEET-knockout Mice

To confirm the complex formation of other proteins with mitoNEET, we performed a Blue Native Polyacrylamide Gel Electrophoresis (BN-PAGE) retardation assay in mitochondrial fractions derived from control mice and mitoNEET-knockout mice. This assay identified the smear in control mice (**Appendix Fig S3, left lane**). In contrast, these bands were almost completely absent in mitoNEET-knockout mice, especially in the range of 50-150 kDa (**Appendix Fig S3, right lane**). This clearly shows the specific formation of complexes with mitoNEET. TfR expression levels in the mitochondria from mitoNEET-knockout mice was lower than from control mice, suggesting that mitoNEET interact with TfR on the OMM *in vivo* (**Fig 3B**).

### mitoNEET Regulates Mitochondrial Iron Contents

The iron contents in isolated mitochondria were significantly higher in mitoNEET-knockout than control mice (**Fig 3C**). In parallel with the iron contents, the levels of mitochondrial ferritin, a mitochondrial iron storage protein, was significantly higher in mitoNEET-knockout than control mice (**Appendix Fig S4A**). However, there was no difference between the two groups in levels of other proteins involved in mitochondrial iron homeostasis, i.e., mitoferin2 (MFRN2, **Appendix Fig S4B**), frataxin (FXN, **Appendix Fig S4C**), adenosine triphosphate (ATP)-binding cassette protein B7 (ABCB7, **Appendix Fig S4D**), and ATP-binding cassette protein B8 (ABCB8, **Appendix Fig S4E**); proteins involved in cytosolic iron homeostasis, i.e., TfR (**Appendix Fig S4F**), divalent metal transporter 1 (DMT1, **Appendix Fig S4G**), and ferroportin (Fpn, **Appendix Fig S4H**); proteins involved in cellular iron homeostasis, i.e., iron regulatory protein 1 (IRP1, **Appendix Fig S4I**) and iron regulatory protein 2 (IRP2, **Appendix Fig S4J**). Heme was not increased in the whole hearts and mitochondria of mitoNEET-knockout mice (**Appendix Fig S4K, S4L**), and proteins involved in heme synthesis also did not differ between the groups (**Appendix Fig S4M, S4N**).

### Deletion of mitoNEET Affects Mitochondrial ROS and Respiration

To assess mitochondrial ROS, we measured the mitochondrial Hydrogen peroxide (H_2_O_2_) release rate from isolated mitochondria during mitochondrial respiration. The mitochondrial H_2_O_2_ release rate during state 3 with glutamate and malate (complex I-linked substrates; GM3) was significantly higher in mitoNEET-knockout than control mice (0.062 ± 0.003 vs. 0.045 ± 0.003 nmol/min/mg mitochondrial protein, P<0.05, **Fig 3D**). After the addition of carbonyl cyanide-p-trifluoromethoxyphenylhydrazone (FCCP), an uncoupler, mitochondrial ROS release during maximal capacity of the electron-transfer system was higher in mitoNEET-knockout than control mice (0.068 ± 0.004 vs. 0.049 ± 0.004 nmol/min/mg mitochondrial protein, P<0.05, **Fig 3D**). In contrast to ROS production, mitochondrial state 3 respiration with glutamate and malate was comparable between the mitoNEET-knockout and control mice (620 ± 120 vs. 712 ± 118 pmol/s/mg mitochondrial protein, **Fig 3E**). There was also no significant difference in mitochondrial maximal capacity between the groups (2331 ± 146 vs. 2689 ± 168 pmol/s/mg mitochondrial protein, **Fig 3E**).

### Deletion of mitoNEET Promotes Mitochondrial and Cardiac Dysfunction by Aging

At the age of 12 months, the iron contents in isolated mitochondria were higher in mitoNEET-knockout than control mice (**Fig 3F**). Mitochondrial H_2_O_2_ release was significantly higher in mitoNEET-knockout mice than control mice both during state 3 respiration (0.080 ± 0.007 vs. 0.049 ± 0.004 nmol/min/mg mitochondrial protein, P<0.05) and maximal capacity (0.084 ± 0.007 vs. 0.056 ± 0.005 nmol/min/mg mitochondrial protein, P<0.05), respectively (**Fig 3G**). Mitochondrial state 3 respiration with glutamate and malate was comparable between mitoNEET-knockout and control mice at 12 months of age (545 ± 52 vs. 571 ± 59 pmol/s/mg mitochondrial protein, **Fig 3H**). In contrast, mitochondrial maximal capacity was significantly lower in mitoNEET-knockout than control mice at 12 months of age (1960 ± 73 vs. 2458 ± 57 pmol/s/mg mitochondrial protein, P<0.05, **Fig 3H**). In short, the mitochondrial reserve capacity, calculated by subtracting state 3 respiration from potential maximum respiratory capacity, was significantly lower in mitoNEET-knockout than control mice (512 ± 83 vs. 1057 ± 48 pmol/s/mg mitochondrial protein, P<0.05, **Fig 3I**).

Cardiac function was evaluated by echocardiography in control and mitoNEET-knockout mice at 12 months of age. Left ventricle (LV) end-diastolic diameter and LV end-systolic diameter, each of which typically dilates in the state of cardiac failure, were significantly higher. As a simple and widely used measure of LV contractility, fractional shortening was significantly lower compared to control mice, suggesting that the heart of 12 months-old mitoNEET-knockout mice had LV dysfunction. LV wall thickness was not different between 12-month-old control and mitoNEET-knockout mice, representing no evidence of cardiac hypertrophy by deletion of mitoNEET (**Fig 4A, 4B**). Histological analysis showed that myocyte cross-sectional area did not differ between groups (**Appendix Fig S5A, S5B**).

**Figure 4.**
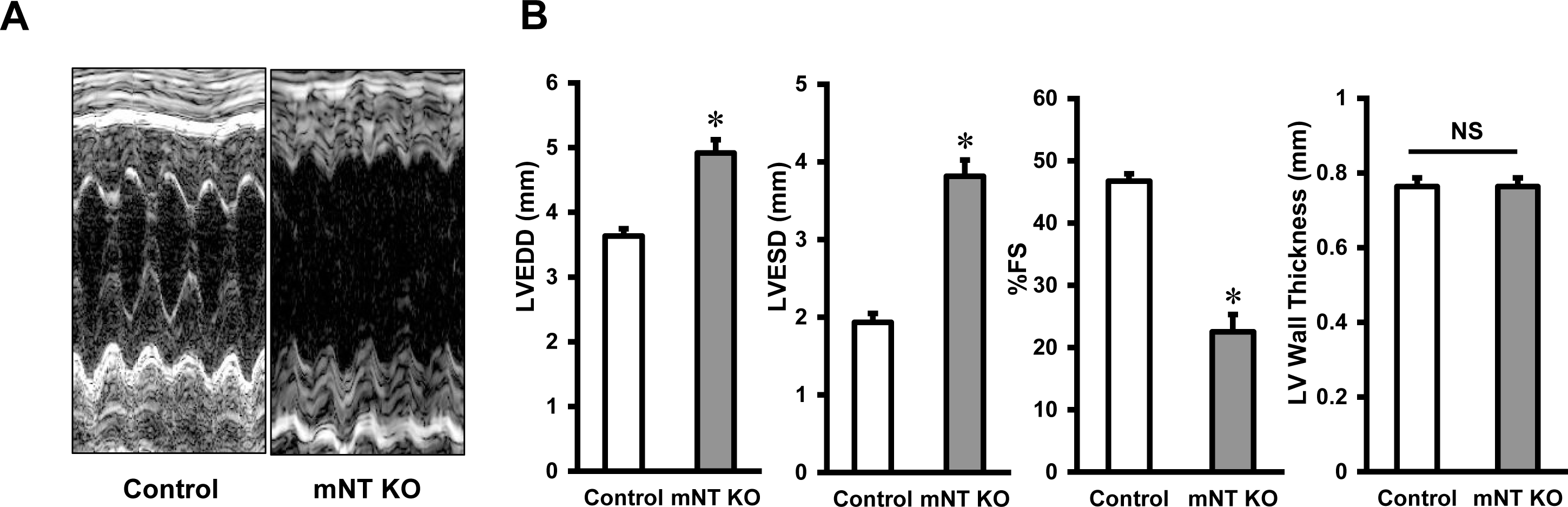
Cardiac Function of 12-month-old mitoNEET-knockout mice assessed by Echocardiography. (A) Representative echocardiography; Control mice(left panel) and mitoNEET-knockout mice (right panel) at the age of 12 months. (B) Summary data of LVEDD (mm), LVESD (mm), %FS, and LV wall thickness in 12-month-old control and mitoNEET-knockout mice. Data are shown as the mean ± SE. *P<0.05 vs. the Control. LV, left ventricle; EDD, end-diastolic diameter; ESD, end-systolic diameter; %FS, percent fractional shortening.

### Aging Decreases mitoNEET Expression and Increases Mitochondrial Iron Contents

Time course of mitoNEET protein expression was examined in the whole heart of C57BL/6J mice at the ages of 3, 6, 9, and 12 months. The expressions of mitoNEET, which were normalized by GAPDH and cytochrome c oxidase IV (COX IV), were significantly lower in the 12-month-old than the 3-month-old mice (**Fig 5A, 5B**). This suggests that mitoNEET protein is decreased in the aged heart independently of mitochondrial quantity. The iron content in isolated mitochondria was higher in 12-month-old than 3-month-old mice (**Fig 5C**). Moreover, the levels of mitoNEET protein tended to be lower in the kidneys of 12-month-old than in those of 3-month-old mice (**Appendix Fig S6**).

**Figure 5.**
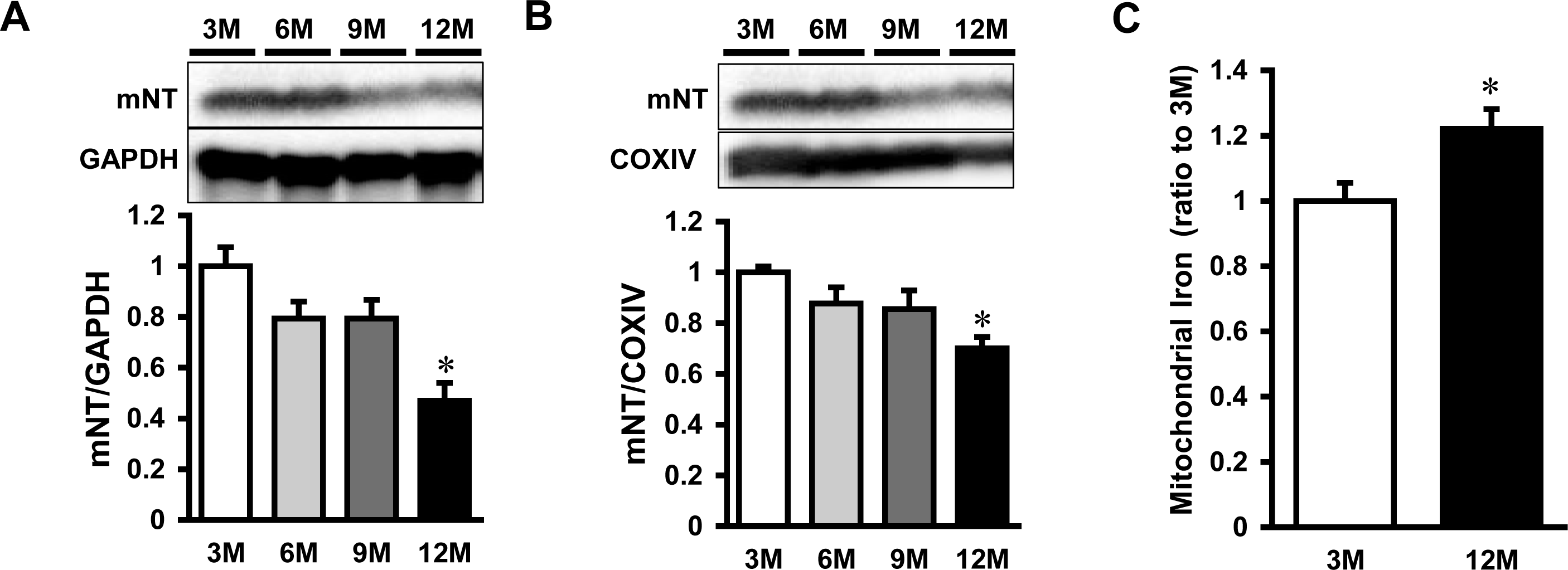
Expression of mitoNEET and Accumulation of Mitochondrial Iron in 12-Month Old C57B6/J Mice. Representative immunoblot and summary data of mitoNEET protein expression normalized to GAPDH (A) and COX IV (B) in the hearts of 3, 6, 9, and 12-month old mice. (C) Levels of mitochondrial iron contents in 12-month old mice relative to 3-month old mice. Data are shown as the mean ± SE. n=5-6. *P<0.05 vs. 3M. M, month; mNT, mitoNEET; GAPDH, glyceraldehyde phosphate dehydrogenase; COX, cytochrome c oxidase.

## Discussion

The main findings of our study were that the endogenous mitoNEET interacted with TfR, a carrier protein for transferrin with iron. Interestingly, mitochondrial iron contents from the hearts of cardiac-specific mitoNEET-knockout mice were higher than control mice. The increase in mitochondrial iron contents corresponded with increased mitochondrial ROS release without affecting mitochondrial respiratory function at the age of 3 months. Therefore, these results suggested that mitoNEET regulated mitochondrial iron homeostasis by protein-protein interaction with TfR.

We focused on a newly discovered novel protein, mitoNEET, located at the OMM (Wiley et al., 2007a). To realize the importance of mitoNEET *in vivo*, we search for clues from the aspect of protein-protein interactions. Namely mitoNEET were selected as target protein for its location of TfR (**Fig 1**). We for the first time clarified that mitoNEET protein interacted with TfR, by using mass spectrometry analysis and immunoprecipitation (**Fig 1B, 1C, 1D, 1E, 2A, 2B, Appendix Table S1, S2**). TfR enters the cellular cytosol in the form of the iron-Tf-TfR complex via endocytosis from the cytoplasmic membrane. To date, TfR has been considered to be in the plasma membrane and endosomes. On the other hand, we also clearly showed that TfR existed with the mitochondria facing the OMM (**Fig 2**), even if we did not probe direct interaction between mitoNEET and TfR. As a previous study said, there are possibility that mitoNEET indirectly interact with TfR via endosomes containging iron-Tf-TfR complex (Das et al., 2016). Because of its location, mitoNEET could play a central role in its complexation with TfR to form TfR-mitoNEET. We propose that mitoNEET may help TfR to primary function as negative regulator of iron inflow into the mitochondria by its interaction with TfR. In other words, that TfR exists on the mitochondria with mitoNEET, not in the membrane or cytosol, may limit iron inflow into the mitochondria, even if the way of its regulation is not elucidated yet.

The relation between mitoNEET protein levels and mitochondrial iron contents does not always reveal the inverse correlation. Because mitochondrial iron concentrations in mammals need to be tightly regulated, iron increase by mitoNEET decrease may be compensated by other known regulatory mechanism in mitochondrial iron homeostasis. Even if that is case, mitoNEET plays a primary role in mitochondrial iron homeostasis. Next, we created mice with cardiac specific deletion of mitoNEET (**Fig 3A, Appendix Fig S2**) and clearly demonstrated that absence of mitoNEET directly increased mitochondrial iron contents (**Fig 3C**). This increase in mitochondrial iron sustained in 12-moths-old mitoNEET-knockout mice as well (**Fig 3F**). Furthermore, there were no differences in the expressions of proteins known to be involved in mitochondrial iron homeostasis, suggesting the existence of an undisclosed pathway of mitochondrial iron homeostasis via mitoNEET. The mitoNEET protein is known to be an acceptor of ISCs (Wiley et al., 2007b). It has also been proposed that the mitoNEET protein transfers iron/sulfur clusters by redox regulation, and donates the clusters and iron ions to mitochondria (Paddock et al., 2007). However, the details of the mechanisms for the acceptance and donation of iron in the mitoNEET protein have not been elucidated.

In the preset study, under the condition of mitochondrial iron overload with the deletion of mitoNEET, the increase in ROS release was detected (**Fig 3D**), measured by using Amplex UltraRed, which was monitored as the fluorescent compound resorufin after converting superoxide anion (O_2_·^−^) into H_2_O_2_. Although we could not clarify the cause-and-effect relationship between iron overload and H_2_O_2_ release, iron plays a crucial role in the redox reaction *in vivo*, and its overload can cause free radical production through many pathways via reduction of O_2_. Moreover, highly reactive ROS are generated by means of the Fenton reaction in the presence of endogenous iron. Therefore, mitochondrial iron overload can easily enhance O_2_·^−^ production via mitochondrial oxidative phosphorylation (OXPHOS), even if overall mitochondrial function is preserved (**Fig 3E**). It was consistent with the overall mitochondrial function and cardiac function in 3-month-old mice (**Fig 3E**). These results suggest that disruption of mitoNEET primarily causes mitochondrial iron overload and enhances ROS production, which secondarily leads to mitochondrial dysfunction. Many previous reports have shown that enhanced ROS production leads to cardiac dysfunction and development of heart failure (HF). We previously reported that an exposure of H_2_O_2_ to cardiac myocytes lead to their injury (Ide, Tsutsui et al., 1999). Furthermore we reported that mitochondria-derived ROS production was increased in the heart from HF model mice (Kinugawa, Tsutsui et al., 2000), and that the treatment with anti-oxidant and overexpression of mitochondrial antioxidant, such as peroxiredoxin-3, improved cardiac function and HF (Matsushima, Ide et al., 2006). Therefore, long-term exposure of ROS over physiological level could lead to cardiac dysfunction. A previous study reported that strong ROS exposure decreased mitochondrial OXPHOS capacity and reserve capacity, resulting in cell death (Dranka, Hill et al., 2010). In contrast, mild ROS exposure decreased mitochondrial reserve capacity, but mitochondrial OXPHOS capacity was almost constant (Dranka et al., 2010). Although the increase in ROS release was the same between 3-month-old mice and 12-month-old mice, the mitochondrial reserve capacity was decreased in 12-month-old mitoNEET-knockout mice, but the mitochondrial OXPHOS capacity was not decreased (**Fig 3G, 3H, 3I**). A previous study showed that deletion of mitoNEET in the liver after high-fat diet feeding increased mitochondrial iron contents and mitochondrial respiration (Kusminski et al., 2012). In contrast, another study reported that the maximal uncoupler-stimulated (state 3u) respiration and complex I-dependent (state 3) respiration were decreased in the isolated mitochondria from the heart of systemic mitoNEET-knockout mice compared to WT mice (Wiley et al., 2007a). This data supports our results. The reason for these discrepancies remains unknown. However, the deletion of mitoNEET can increase mitochondrial iron contents, and decrease mitochondrial respiration at least in the heart. In accordance with mitochondrial reserve capacity, chronic ROS exposure caused cardiac dysfunction (**Fig 4**).

Finally, we showed that mitochondrial iron was increased by aging in association with a decrease in mitoNEET protein (**Fig 5**). This suggests that the mitoNEET-iron relationship is an important physiological and universal phenomenon, but not one specific to cardiac diseases. Moreover, in the kidney, the decrease of mitoNEET expression in aging mice were observed as well, this result also support the importance of mitoNEET (**Appendix Fig S6**).

Mitochondrial iron has been reported to play an important role in some cardiac diseases. Doxorubicin, an anthracycline antibiotic, is one of the most widely used agents as chemotherapy for hematological malignancies and solid tumor, which often induces cardiac dysfunction and HF, i.e. doxorubicin-induced cardiomyopathy (Minotti, Menna et al., 2004). Previous clinical study showed that levels of mitochondrial iron were significantly higher in the explanted heart from patients with doxorubicin-induced cardiomyopathy than in the heart from normal subjects and patients with other cardiomyopathies (Ichikawa, Ghanefar et al., 2014). Increased ROS caused by iron overload are thought to be its major cause. Thus, mitochondrial iron homeostasis plays an important role on cardiac function. Its regulation has been controlled by several proteins in mitochondria. FXN, located in mitochondria matrix, plays a role in synthesis of iron/sulfur cluster (Vaubel & Isaya, 2013). Friedreich’s ataxia (FRDA), a human genetic disease caused by GAA triplet expansion of FXN gene, leads to hypertrophic cardiomyopathy, as well as neurodegeneration (Isaya, 2014). Mouse model of FXN deletion in the heart, which mimic FRDA cardiomyopathy, by gene therapy with adeno-associated virus rh10 vector expressing human FXN, prevents cardiomyopathy progressing and also reverse cardiomyopathy with HF (Perdomini, Belbellaa et al., 2014). In addition, Mice with knockout of ABCB8, which is localized in the inner mitochondrial membrane and functions as iron exporter from the mitochondrial matrix, showed the progression of LV dysfunction in association with iron accumulation and an increase in ROS (Ichikawa et al., 2012). In contrast, iron within mitochondria is used for the synthase of iron/sulfur clusters and heme. Therefore, mitochondrial iron needs to be finely controlled under every kind of conditions around mitochondria. In the present study, we clearly demonstrated that mitoNEET was the regulator of iron homeostasis.

In summary, we show that the increase in mitochondrial iron contents corresponded with increased mitochondrial ROS release in the hearts of cardiac-specific mitoNEET-knockout mice. The endogenous mitoNEET interacted with TfR, suggesting that mitoNEET regulated mitochondrial iron homeostasis by protein complex formation. These results suggest that the regulation of mitochondrial iron would be a potential target for the treatment of cardiac diseases and other conditions.

## Methods

All experimental procedures and methods of animal care were approved by the Institutional Animal Care and Use Committee of National University Corporation Hokkaido University (Permit Number: 16-0101) and also conformed to the Guide for the Care and Use of Laboratory Animals published by the US National Institutes of Health.

### Immunoblot

Samples, 10-20 μg of total protein from heart tissues, were separated by SDS-PAGE and transferred to a polyvinylidene fluoride (PVDF) membrane (Bio-Rad, USA). The membrane was blocked for 1 h at room temperature in TBS-T buffer (Tris buffered saline containing 0.1% Tween 20) containing 3% BSA or milk, and was incubated with the primary antibodies at a dilution of 1:1000 overnight at 4°C. After 3 washings with TBS-T, the membrane was incubated with a horseradish peroxide-conjugated secondary antibody at a dilution 1:5000 for 1 h at room temperature. After washing, the membrane was developed with ECL or ECL Prime Reagent (GE, USA) and then processed for detection with ChemiDoc XRS+ (Bio-Rad, USA). The density of the signals of bands was quantified with Image J (NIH) software.

### Silver-intensified immunogold method for electron microscopy

Cultured C2C12 cells were fixed for 2 hr with 4% paraformaldehyde in 0.1 M phosphate buffer, pH 7.3. After pretreatment in normal donkey serum for 30 min, they were incubated with the mouse anti-TfR monoclonal antibody (1: 2000 in dilution, Thermo Fisher) at 4oC overnight, and subsequently reacted at 4oC overnight with goat anti-mouse IgG covalently linked with 1-nm gold particles (1: 400 in dilution; Nanoprobes, Yaphank, NY). Following silver enhancement using a kit (HQ silver; Nanoprobes), the samples were osmificated, dehydrated, and directly embedded in Epon (Nisshin EM, Tokyo, Japan). Ultrathin sections were prepared and stained with both uranyl acetate and lead citrate for observation under an electron microscope (H-7100; Hitachi, Tokyo, Japan).

### Immunoprecipitation of Endogenous Proteins

To analyze the interaction between endogenous mitoNEET and other proteins, including TfR, the whole cell lysates from mouse heart were solubilized with lysis buffer (Cell Signaling Technology, USA). The lysates were centrifuged at 15,000 rpm for 20 min at 4°C, and then the supernatant fluid was collected. The lysates were mixed with an anti-TfR Ab (Abcam, UK) and normal rabbit IgG as a control, and incubated overnight at 4°C. After the addition of protein-A (rProtein A Sepharose Fast Flow, GE, USA), the lysates were incubated for 4 h at 4°C. Protein-A was washed with buffer (150 mM NaCl, 50 mM Tris-HCl, 0.5% Triton-X) and then mixed with 2 ×; SDS sample buffer with boiling for 5 min at 95°C.

### Cell Culture

The mouse C2C12 myoblast cell line was purchased from the American Type Culture Collection (Manassas, USA). Mouse C2C12 myoblasts were seeded at a concentration of 4×10^5^ cells/ml in 6-well culture plates. Differentiation of C2C12 myoblasts into myotubes was induced by medium containing 2% horse serum for 24 h, as previously described(Fukushima, Kinugawa et al., 2014).

### Iron Addition or Reduction for Cells

When mouse C2C12 cells reached confluence, 20 μM ferric ammonium citrate (FAC) (Sigma) in addition to iron, and desferioxamine (DFO) (Sigma) in reduction of iron, were added 24h.

### Preparation of Isolated Mitochondria

Heart tissues were quickly harvested, and mitochondria were isolated (Tonkonogi & Sahlin, 1997). Briefly, heart tissues were minced on ice, and incubated with mitochondrial isolation buffer containing 0.1 mg/ml proteinase (Sigma-Aldrich, USA) for 2 min. The heart tissue was gently homogenized with six strokes using a motor-driven Teflon pestle in glass chamber. The homogenate was centrifuged at 750 *g* for 10 min. The supernatant was centrifuged at 10,000 *g* for 10 min, and the pellet was washed and centrifuged at 7,000 *g* for 3 min. The final pellet was suspended in suspension buffer (containing 225 mmol/l mannitol, 75 mmol/l sucrose, 10 mmol/l Tris, and 0.1 mmol/l EDTA; pH 7.4). Finally, the mitochondrial protein concentration was measured using a BCA assay.

### Digitonin treatment

Digitonin was dissolved in DMSO. To extract cholesterol from the OMM, isolated mitochondria were incubated for 5 min at room temperature with 0.03 mg/ml digitonin. To solubilize isolated mitochondria, digitonin at the indicated final concentrations was added to 100 µl of 1.2 mg/ml mitochondria in mitochondrial suspension buffer (Tris-HCl (pH 7.4), 75 mM sucrose, and 225 mM mannitol). After 5 min incubation at room temperature and centrifugation at 10,000 *g*, the pellet was suspended in 100 µl of 2 X sample buffer. Equivalent portions of the supernatant and pellet were subjected to SDS-PAGE and evaluated by immunoblot (Arasaki et al., 2015).

### Immunohistochemical Staining

For immunohistochemical staining, the mouse C2C12 cells were plated onto cover-glasses coated with 50 µg/ml fibronectin. After 24-hr incubation at 37°C, the cells were fixed with 2% formaldehyde in phosphate-buffered saline (PBS) at 37°C for 10 min, followed by ice-cold methanol for a 5-min fixation at −20°C. After blocking with 1% bovine serum albumin (BSA) in PBS for 30 min, the samples were incubated with antibodies against mitoNEET and TfR in 1% BSA/PBS overnight at 4°C. After being rinsed with blocking buffer, the samples were incubated with secondary antibodies conjugated with Alexa Fluor Plus 488 (against rabbit IgG) and Alexa Fluor Plus 555 (against mouse IgG) at a dilution of 1:1000 for 30 min at room temperature.

Fluorescence images were obtained with a structured illumination microscopy (N-SIM, Nikon, Japan).

### Experimental Animals

All mice were bred in a pathogen-free environment and housed in an animal room under controlled condition on a 12 h: 12 h light/dark cycle at a temperature of 23°C to 25°C.

C57BL/6J mice (CLEA Japan Inc.) were bred under controlled conditions and euthanized under deep anesthesia with tribromoethanol-amylene hydrate (Avertin; 250 mg/kg body weight, i.p.) (Kinugawa et al., 2000) at the age of 3, 6, 9 or 12 months (n=6, each). The heart and kidney were excised.

Mice with disruption of cardiac-specific mitoNEET were newly generated (**Fig S1**). Briefly, the conditional mitoNEET allele was created using clones isolated by restriction mapping with a genome library, and a C57BL/6 genetic background clone was used to construct the targeting vector. A neo cassette was inserted downstream from exon 2 and was flanked by FRT sites for later excision by FLP recombinase. The lox-P sites were inserted upstream from exon 2 and downstream from the neo cassette. The targeting vector was transfected by electroporation of embryonic stem cells. After selection, the surviving clones were expanded for polymerase chain reaction analysis to identify recombinant embryonic stem clones. Targeted embryonic stem cells were microinjected into C57BL/6 blastocysts, and chimera mice were mated with wild type C57BL/6 homozygous FLP mice to remove the neo cassette. Heterozygous mice with neo deletion and confirmed lox-P sites were crossed with C57/BL6 mice to obtain heterozygous mice. Finally, mitoNEET flox/flox mice were also crossed with αMHC-Cre mice to obtain cardiac-specific deletion of mitoNEET (mitoNEET-knockout mice). mitoNEET flox/flox mice were used as control mice. Most of experiments were performed at the age of 3 months, with others performed at the age of 12 months.

### Measurement of Mitochondrial Iron Contents

Isolated mitochondria, collected by using a Mitochondrial Isolation Kit for Tissue (Pierce, USA), were diluted with EDTA-free buffer after sonication.

Mitochondrial “non-heme” iron contents were measured by using a commercial Iron Assay Kit (BioAssay Systems, USA), which directly detect total iron in the sample, according to the manufacturer’s protocol(Kusminski et al., 2012). Isolated mitochondria with sonication were diluted with EDTA-free buffer, and normalized to the mitochondria concentration of each sample.

### Mitochondrial OXPHOS Capacity and ROS Release

The mitochondrial respiratory capacity was measured in isolated mitochondria at 37°C with a high-resolution respirometer (Oxygraph-2k, Oroboros Instruments, Austria) (Christiansen, Dela et al., 2015, Takada, Masaki et al., 2016). H_2_O_2_ release from isolated mitochondria was measured at 37°C by spectrofluorometry (O2k-Fluorescence LED2-Module, Oroboros Instruments, Austria). The release of H_2_O_2_ that we measured reflected intrinsic O_2_·^−^ and H_2_O_2_ release in the mitochondria under the presence of superoxide dismutase (SOD) (Hey-Mogensen, Hojlund et al., 2010). O_2_·^−^, unstable substance, needs to be converted into H_2_O_2_, relatively stable one, to evaluate ROS production. H_2_O_2_ reacts with Amplex UltraRed (Life Technologies, USA) in an equal amount of stoichiometry catalyzed by horseradish peroxidase, which yields the fluorescent compound resorufin (excitation: 560 nm; emission: 590 nm). Resorufin was monitored throughout the experiment. Beforehand five different concentrations of H_2_O_2_ were added to establish a standard curve in advance. The H_2_O_2_ release rate from isolated mitochondria is expressed as nanomoles per minute per milligram of mitochondrial protein.

After the addition of isolated mitochondria (approximately 100-200 µg) to the chamber in the respirometer filled with 2 ml of MiR05 medium with 5 U/ml SOD, 25 µmol/l Amplex ultrared, and 1 U/ml horseradish peroxidase, substrates, adenosine diphosphate (ADP), and inhibitors were added in the following order: 1) glutamate 10 mmol/l + malate 2 mmol/l (complex I-linked substrates), 2) ADP 10 mmol/l, 3) succinate 10 mmol/l (a complex II-linked substrate), 4) oligomycin 2.5 μmol/l, 5) FCCP 1 μmol/l, 6) rotenone 0.5 μmol/l, 7) antimycin a 2.5 μmol/l. O_2_ consumption rates, i.e., respiratory rates, were expressed as O_2_ flux normalized to mitochondrial protein concentration (µg/µl). Datlab software (Oroboros Instruments) was used for data acquisition and data analysis.

### Echocardiographic Measurements

Echocardiographic measurements were performed under light anesthesia with tribromoethanol/amylene hydrate (avertin; 2.5% wt/vol, 8 μL/g i.p.). Two-dimensional parasternal short-axis views were obtained at the levels of the papillary muscles. Two-dimensional targeted M-mode tracings were recorded at a paper speed of 50 mm/sec.

### Statistical Analysis

Data are expressed as the mean ± the standard error of the mean (SE). Student’s unpaired *t*-tests were performed to compare means between two independent groups. One-way ANOVA followed by the Tukey’s test was performed for multiple-group comparisons of means. Values of P<0.05 were considered statistically significant.

**Figure 6.**
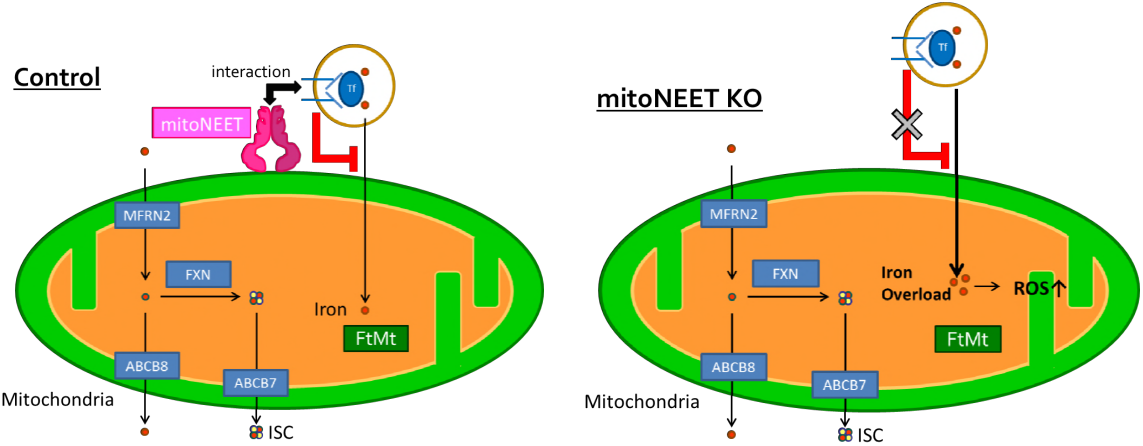
A schematic mechanism of iron overload in mitoNEET-knockout mice. The interaction between mitoNEET and transferrin receptor causes mitoNEET to be colocalize with transferrin receptor, which in control limits inflow of iron. In contrast, the absence of mitoNEET in mitoNEET-knockout mice increases inflow of iron, for transferrin receptor was separated from mitochondrial outer membrane, leading to mitochondrial iron overload and consequently enhancing mitochondrial ROS production. ROS, reactive oxygen species.

## Acknowledgements

We thank Yuki Kimura, Noriko Ikeda and Miwako Yamane for their technical assistance. This work was supported in part by grants from Japanese Grant-In-Aid for Scientific Research (JP17K15979 (T.F.), JP17K10137 (A.F.), JP17H04758 (S.T.), 18K08022 (T.Y.), 18H03187 (S.K.), 26350879 (S.K.), 15H04815 (H.T.)), the Japanese Association of Cardiac Rehabilitation (T.F.), Hokkaido Heart Association Grant for Research (T.F., S.T.), Suhara Memorial Foundation (S.K), the Mochida Memorial Foundation for Medical and Pharmaceutical Research (T.Y.), the Nakatomi Foundation (T.Y.), the Japan Foundation for Applied Enzymology (S.T.), Northern Advancement Center for Science & Technology (S.T.), Japan Heart Foundation & Astellas Grant for Research on Atherosclerosis Update (S.T.), MSD Life Science Foundation (S.T.), Uehara Memorial Foundation (S.T.), Cardiovascular Research Fund, Tokyo, Japan (S.T.), SENSHIN Medical Research Foundation (S.T.), the Nakatomi Foundation (S.T.), Japan Heart Foundation (S.T.), the Sasakawa Scientific Research Grant from The Japan Science Society (S.T.).

## Author contributions

T.F., S.T. and S.K. designed the study. M.W., H.T., S.H., M.M. and K.I.N. performed Mass Spectrometry Analysis. N.K., Y.O., H.S. performed IHC analysis. J.N-K and T.I performed immune electron microscopy. T.F., S.T., S.M., A.F., M.T., and J.M. performed the other experiments. T.F. and S.K. wrote the manuscript with help from T.Y., S.M., and H.T.

## Conflict of interset

The authors have no conflicts of interest to disclose.

## Appendix

### Methods

#### Mass Spectrometry Analysis

HEK 293 cells were transfected with or without expressed 3 ×; FLAG-mitoNEET under the control of a cytomegalovirus constitutive promoter in the pcDNA3 expression vector. The cells were lysed, and the lysate was centrifuged. The resultant supernatant was incubated at 4°C with the antibody to FLAG (M2, Sigma-Aldrich, USA) immobilized on protein-A (rProtein A Sepharose Fast Flow, GE, USA), and the beads were washed with the lysis buffer. The immunoprecipitated proteins were eluted with the 3 ×; FLAG peptide (Sigma-Aldrich, USA).

The proteins collected proteins by immunoprecipitation were separated by SDS-PAGE gel and stained with silver staining. These silver-stained bands were excised from the gels. The proteins therein were subjected to in-gel reduction, S-carboxyamidomethylation and digestion with sequence-grade trypsin (Promega, Fitchburg). The resultant peptides were analyzed by LCESI-MS/MS (LCQ DECA and LTQ XL; Thermo Fisher Scientific, USA). The data were analyzed using Mascot software (Matrix Science, USA) (Yamamoto, Takeya et al., 2013).

#### Genotyping for mitoNEET KO Mice

Genotyping of mitoNEET KO mice was performed by PCR with DNA extracted from the tail. To detect the Cre recombinase, the following primers were used: 5’-CTGAAAAGTTAACCAGGTGAGAATG -3’ (forward) and 5’-AGGTAGTTATTCGGATCATCAGCTA -3’ (reverse). To distinguish mitoNEET flox/flox or wild type, the following primers were used: 5’-TCTAAAATGTACAGCAGCCATGAAG -3’ (forward) and 5’-ACCAAGATACTTAGCGGTAGAAGTG -3’ (reverse). The protocol of PCR amplification was as follows: 35 cycles of 10 sec at 98 °C, 5 sec at 65 °C, and 120 sec at 72 °C; followed by 35 cycles of 10 sec at 98 °C, 5 sec at 66 °C, and 60 sec at 72 °C, respectively.

#### Quantitative Reverse Transcriptase PCR

Total RNA was extracted from heart tissues with QuickGene-810 (FujiFilm, Japan) according to the manufacturer’s instructions. The total RNA concentration and purity were assessed by measuring the optical density (230, 260, and 280 nm) with a Nanodrop 1000 Spectrophotometer (Thermo Fisher Scientific, USA). cDNA was synthesized with a high capacity cDNA reverse transcription kit (Applied Biosystems, USA). Reverse transcription was performed for 10 min at 25°C, for 120 min at 37°C, for 5 sec at 85°C, and then solution was cooled at 4°C. TaqMan quantitative realtime PCR was performed with the 7300 real-time PCR system (Applied Biosystems) to amplify samples for Cisd1 (Mm01172641_g1) cDNA in the heart. After 2 min at 50°C and 10 min at 95°C, the PCR amplification was performed for 40 cycles of 15 sec at 95°C and 1 min at 60°C. GAPDH was used as an internal control. Data were analyzed using a comparative 2^−ΔΔCT^ method.

#### Generation of mitoNEET Antibody

The peptide antigen, a C-terminal fragment of mitoNEET, was chemically synthesized. Samples using the peptide were injected into 4-month-old female New Zealand white rabbits. After multiple immunizations, the blood samples were collected from the ear vein of the rabbits. The generation of antibodies was examined by immunoblot, and levels of anti-peptide antibody were determined with the conventional ELISA method.

#### Blue Native Page Electrophoresis (BN-PAGE)

BN-PAGE was performed as previously described (Wittig, Braun et al., 2006). The proteins from the whole heart were extracted with 5% digitonin (Invitrogen) (protein : detergent ratio of 1 : 10) and 4 ×; buffer on ice for 30 min. After centrifugation at 10,000 g for 10 min at 4 °C, the supernatants were collected. The remaining lysate was combined with Coomasie blue G-250 dye (Invitrogen) (protein : detergent ratio of 1 : 10) and added to 3-12% NativePAGETM Novex Bis-Tris Gel (Invitrogen), then separated electrophoretically by SDS-PAGE using an Anode and Cathode buffer (Invitrogen) at 10 mA for 1 h and at 150V for 2 h on ice. The protein complex in the samples after SDS-PAGE was denatured by denaturing buffer (in mmol; Tris 20, glycine 200, 1% SDS). Then the gels were transferred by electroblotting to PVDF membranes (Bio-Rad) using transfer buffer at 30 V for 3 h.

#### Measurement of Heme

About 2-3 mg of heart tissue was homogenized in 1% Triton-X100 in Tris-buffered saline and centrifuged at 5,000 *g* for 10 min. The supernatants were collected, the lysate for total heme measurement was prepared and then the protein concentration was quantified by BCA assay. The lysate for mitochondrial heme measurement was prepared using a Mitochondrial Isolation Kit for Tissue (Pierce, USA).

Levels of total and mitochondrial heme were quantified as previously described (Khechaduri, Bayeva et al., 2013). Briefly, equal amounts of total or mitochondrial proteins were mixed with 2 M oxalic acid and boiled to 95°C for 30 min. After centrifugation at 1,000 *g* for 10 min at 4°C, the supernatants containing release iron from heme and generated fluorescent PPIX were collected. The fluorescence of the supernatant was assessed at 405 nm/600 nm on a Spectra Max Gemini fluorescence microplate reader, which was normalized to the protein concentration of each sample.

#### Organ Histology

For histological analysis, tissue was fixed in 10% formaldehyde, cut into three transverse sections; apex, middle ring, and base, then stained with hematoxylin-eosin. Myocyte cross-sectional area was determined as described previously(Kinugawa et al., 2000).

### Figure Legends

**Appendix Figure S1.**
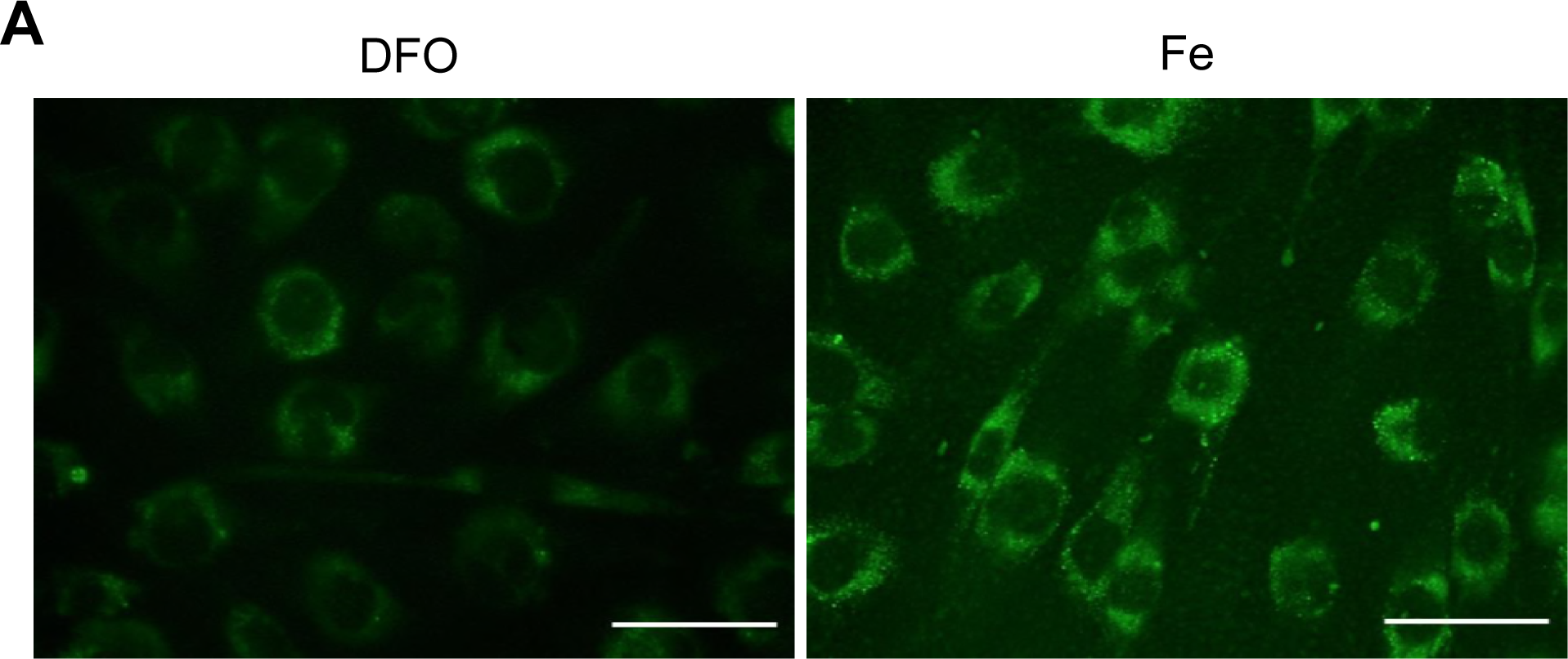
Mitochondrial Iron Contents Influenced by Iron or DFO. Representative images obtained from super-resolution microscopy by using Mito-ferroGreen (Dojindo, Japan). Bar, 50 μm. DFO, desferioxamine; Fe, the addition of ferric ammonium citrate.

**Appendix Figure S2.**
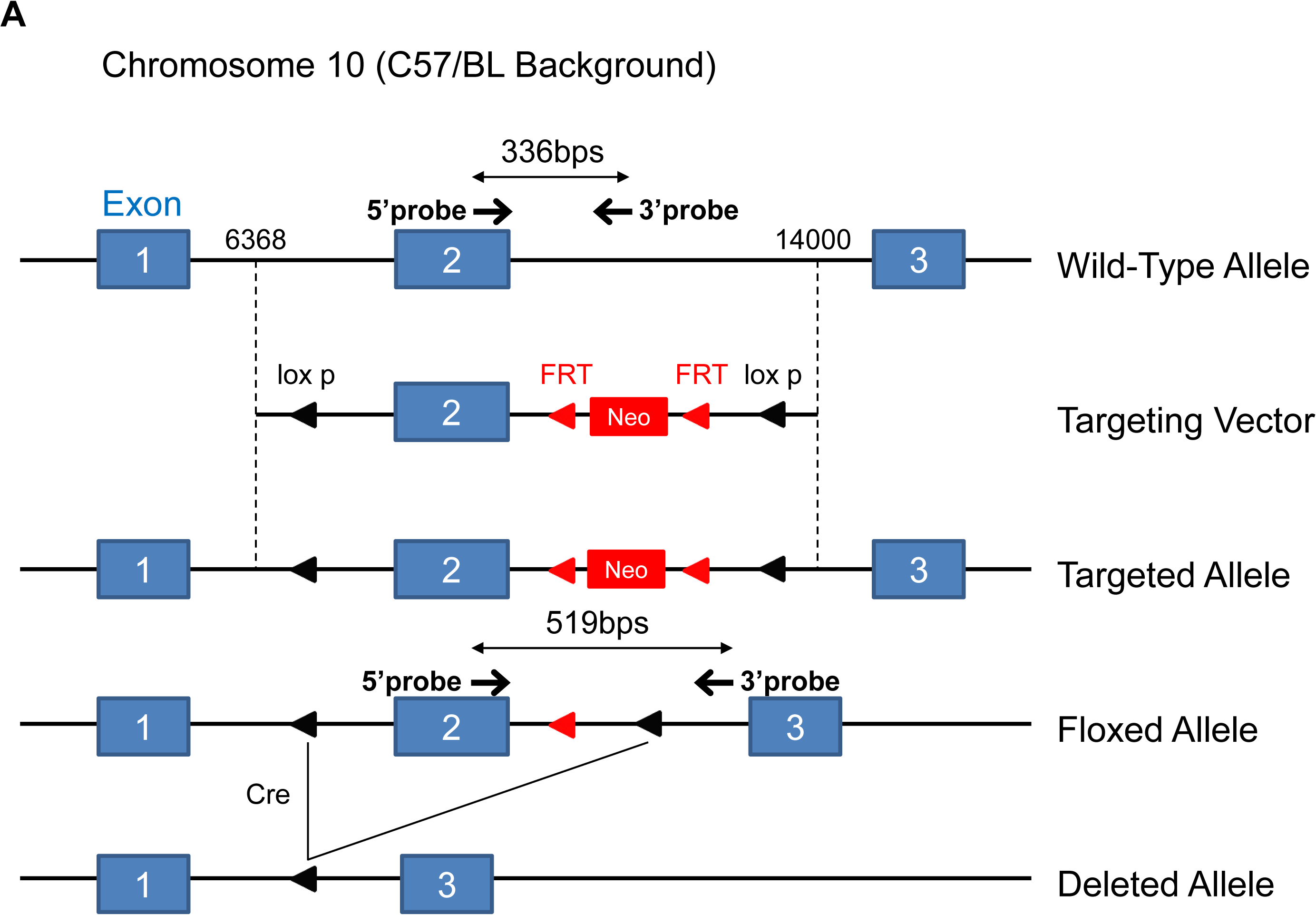
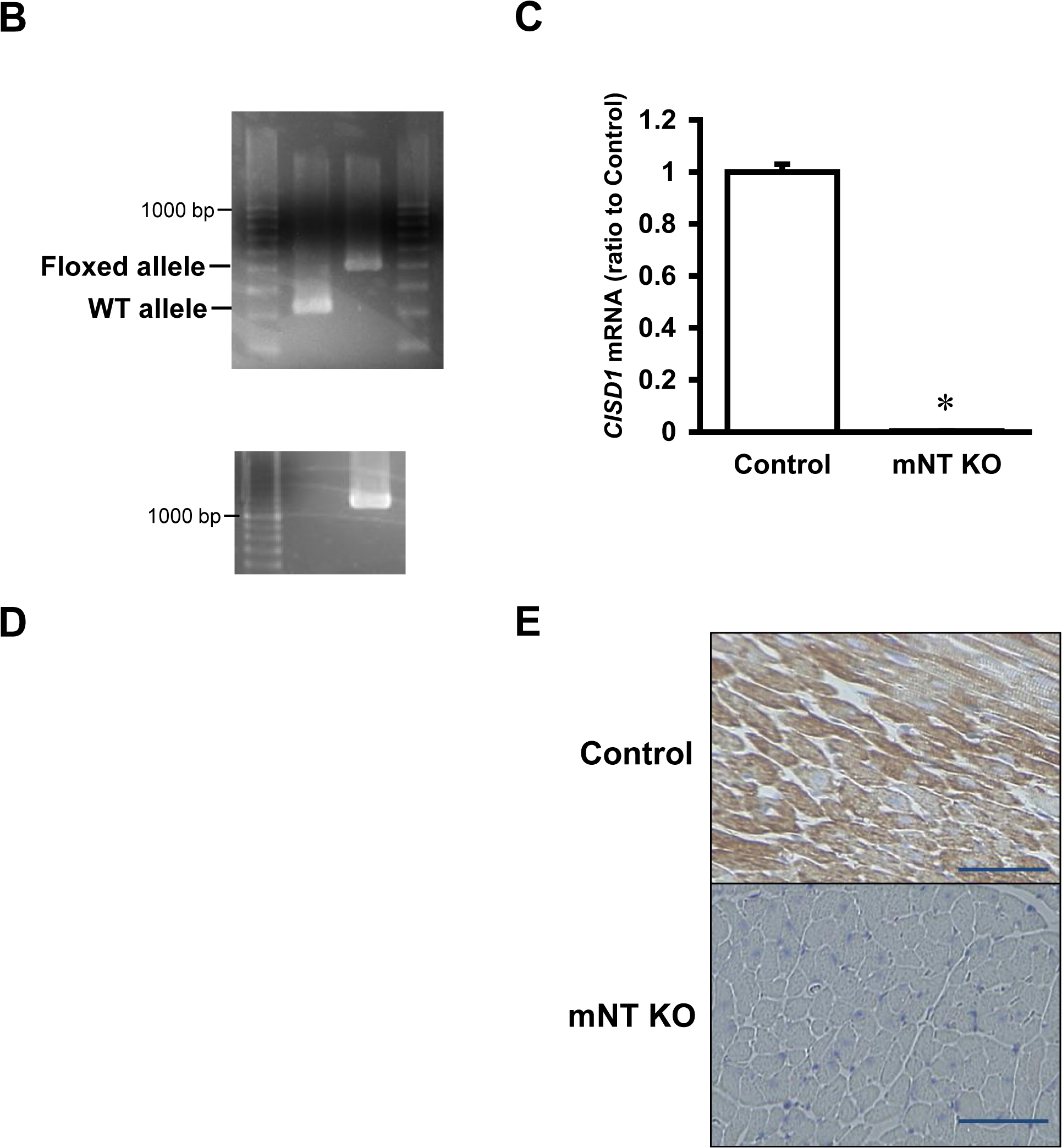
Generation of mitoNEET KO Mice. (A) Design of the mitoNEET targeting construct and the genomic structure of the mitoNEET. LoxP sites were inserted to delete the entire exon 2, resulting in early termination and truncation of the C-terminal region of mitoNEET. This resulted in the complete destruction of mitoNEET function. The indicated primers were used for detecting mitoNEET flox allele. (B) WT and mitoNEET floxed alleles were distinguished by polymerase chain reaction (PCR) analysis. Genomic PCR confirmed the mitoNEET floxed alleles and Cre allele in mitoNEET-knockout mice. (C) Quantitative analysis of the gene expression of *CISD1* in the heart (n=10-11). Data are shown as the mean ± SE. *P<0.05 vs. Control. (D) Representative immunoblot by Tris-Tricine SDS-PAGE of the lysate from control (Ctrl) and mitoNEET-knockout mice (KO), and the peptide of the mitoNEET fragment as positive control (PC). The black arrow (about 14 kDa) indicates mitoNEET, and the white arrow (below 2 kDa) indicates mitoNEET fragment as PC. (E) Representative positive immunostaining for mitoNEET in myocardial sections. Upper Panel: Control; Lower Panel: mNT KO. Scale Bar, 50 μm. WT, wild-type; mNT KO, mitoNEET-knockout; *CISD1*, CDGSH iron sulfur domain 1;CZ, cruz marker; PC, positive control; Ctrl, control; CM, color marker; GAPDH, glyceraldehyde phosphate dehydrogenase.

**Appendix Figure S3.**
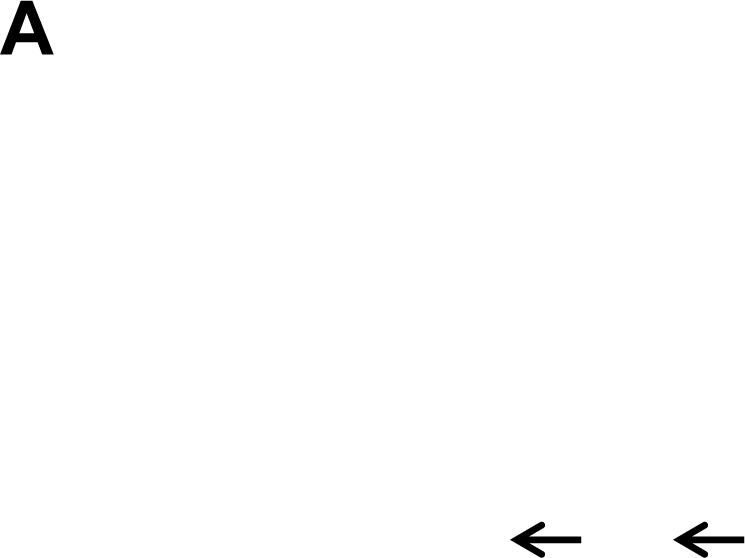
Complex of mitoNEET in mitoNEET-knockout mice. (A) Representative BN-PAGE retardation assays performed on the whole cell lysate from the heart of control mice or mitoNEET-knockout mice. Black arrows (about 120 kDa) indicate the complex consisting of mitoNEET and TfR. IB, immunoblot; IP, immunoprecipitation; KO, knockout.

**Appendix Figure S4.**
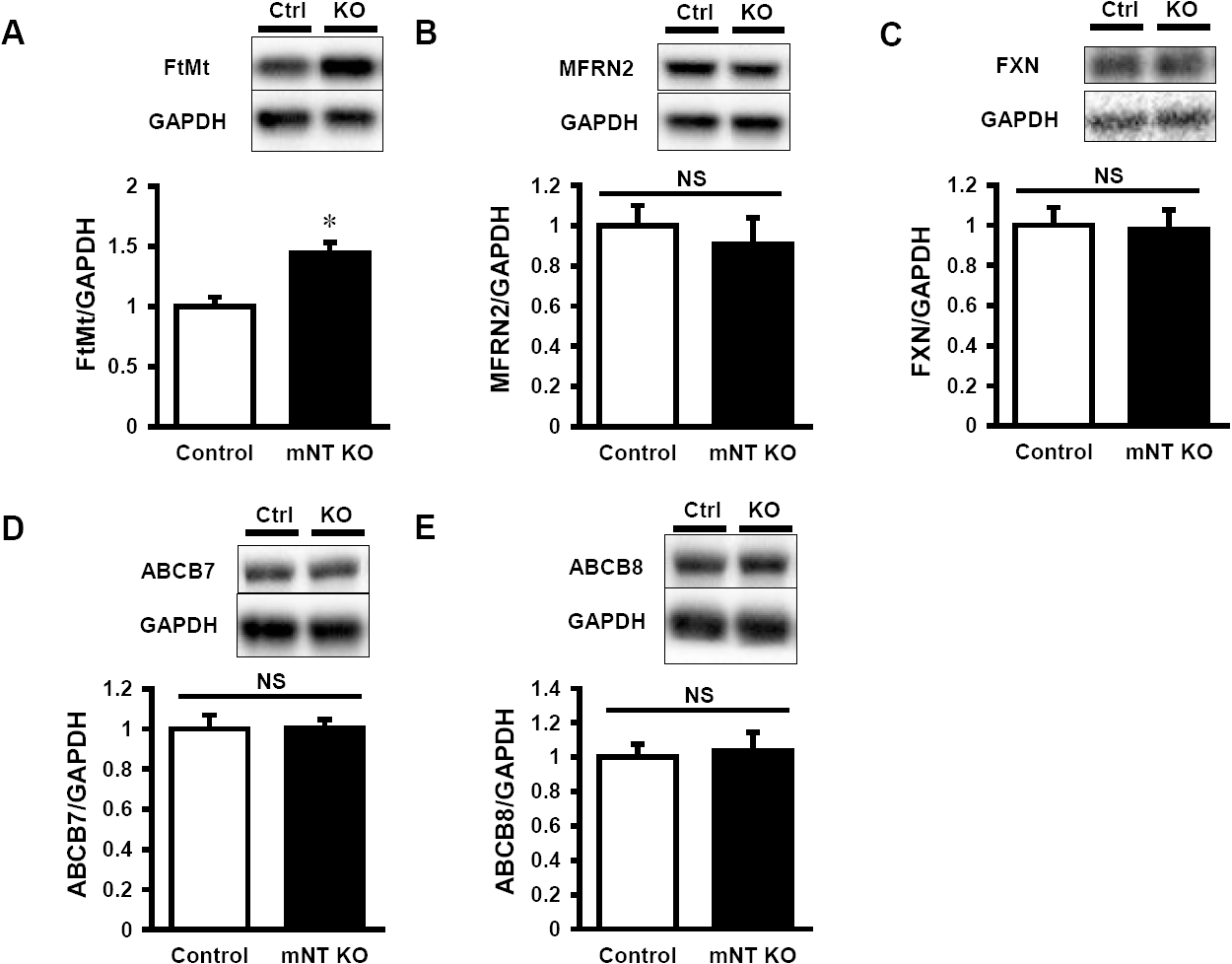
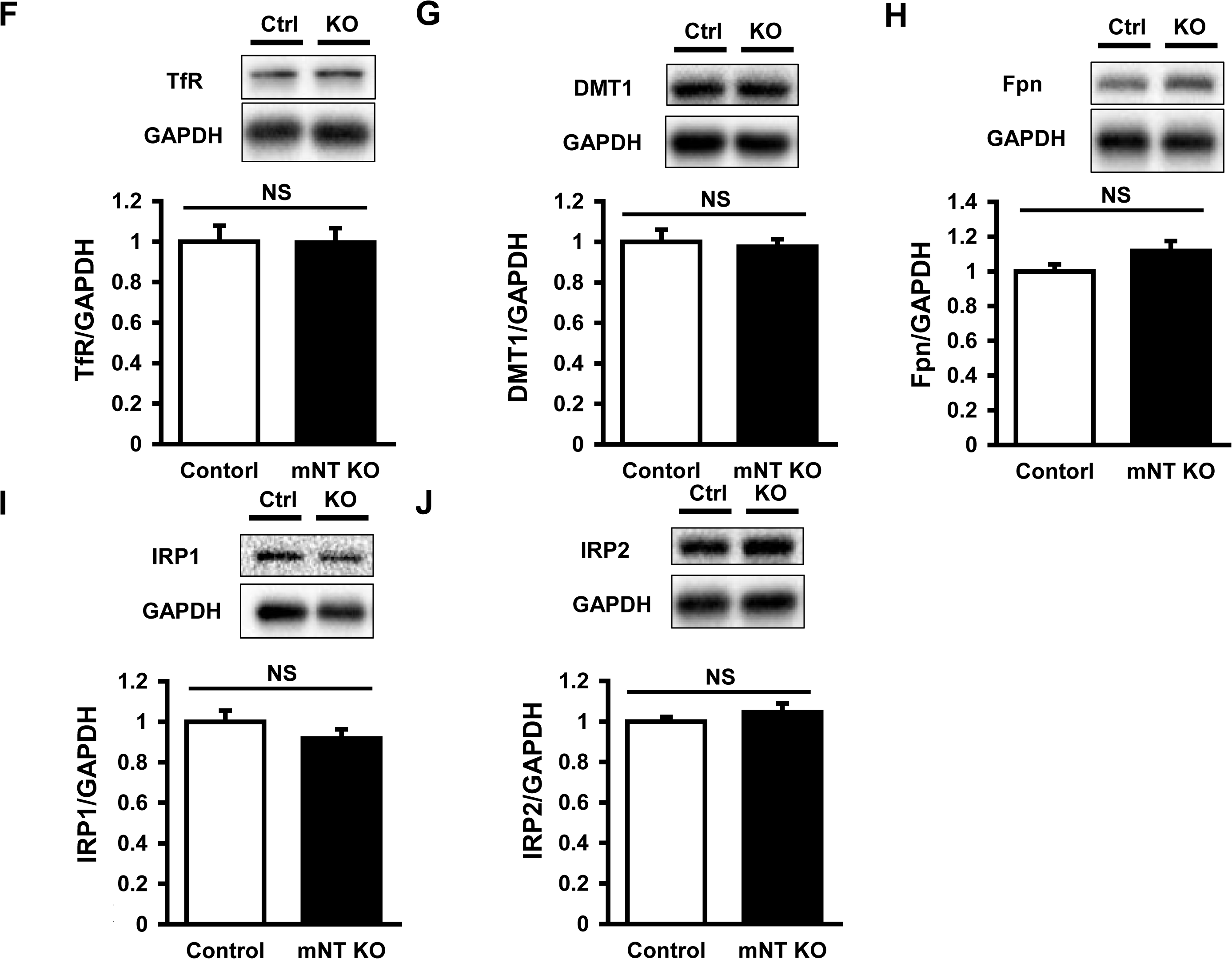
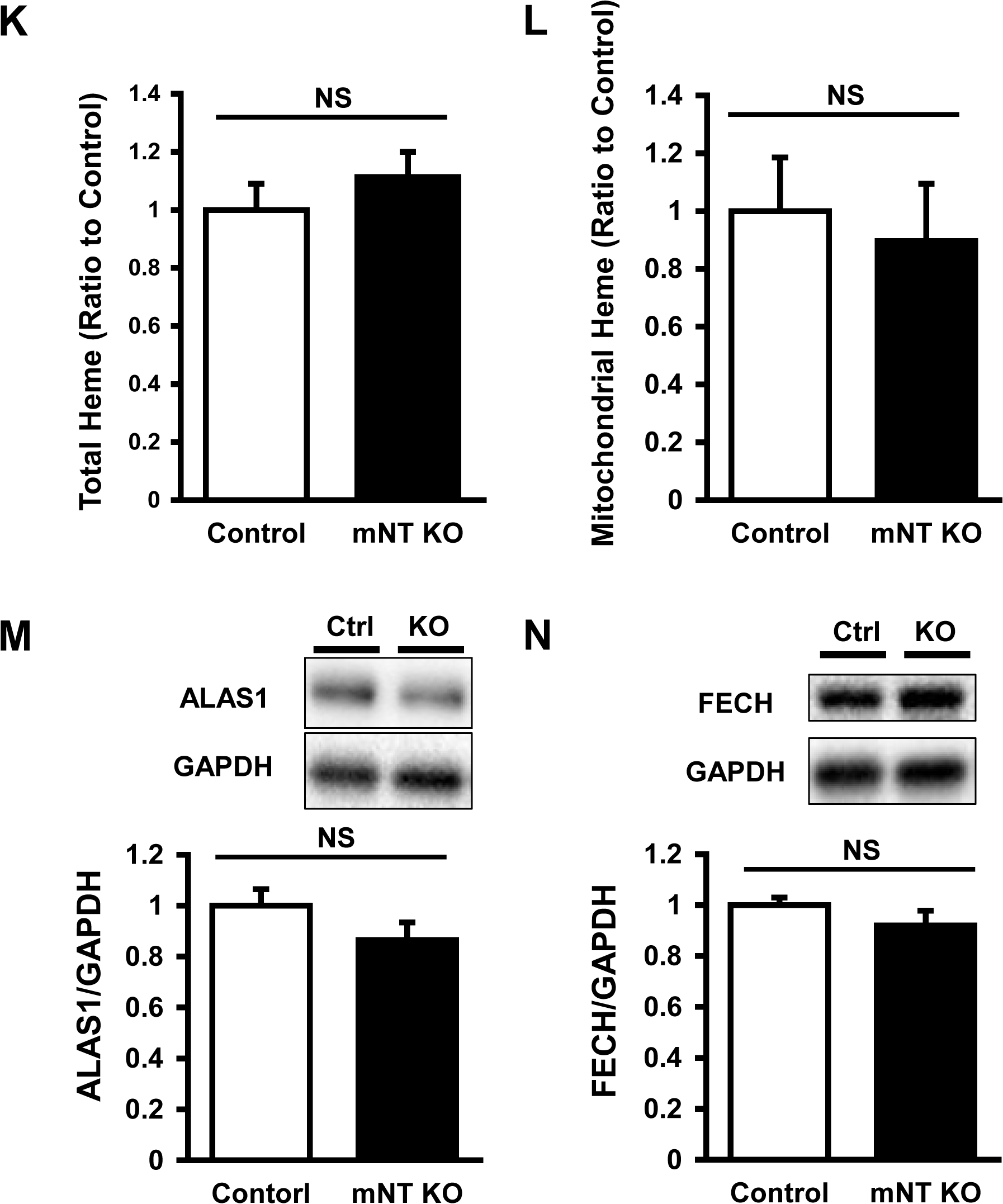
Proteins Related to Iron Homeostasis and Levels of Heme and Expression of Proteins Related to Heme Synthesis in the Heart of 3 Months-Old mitoNEET-knockout mice. Representative immunoblot and summary data of FtMt (A), MFRN2 (B), FXN (C), ABCB7 (D), ABCB8 (E), TfR (F), DMT1 (G), Fpn (H), IRP1 (I), and IRP2 (J) protein expressions normalized to GAPDH in the heart from control and mitoNEET KO mice (n=10-11). Levels of total heme (K) and mitochondrial heme (L) in mitoNEET KO mice relative to control mice (n=7-9). Representative immunoblot and summary data of ALAS1 (M) and FECH (N) protein expressions normalized to GAPDH in the heart from control and mitoNEET KO mice (n=10-11). Data are shown as the mean ± SE. *P<0.05 vs. Control. Ctrl, control; NS, not significant; FtMt, mitochondrial ferritin; MFRN2, mitoferin2; FXN, frataxin; ABCB7, ATP-binding cassette protein B7; ABCB8, ATP-binding cassette protein B8; TfR, transferrin receptor; DMT1, divalent metal transporter 1; Fpn, ferroportin; IRP1, iron regulatory protein 1; IRP2, iron regulatory protein 2; ALAS1, 5’-aminolevulinate synthase 1; FECH, ferrochelatase; GAPDH, glyceraldehyde phosphate dehydrogenase.

**Appendix Figure S5.**
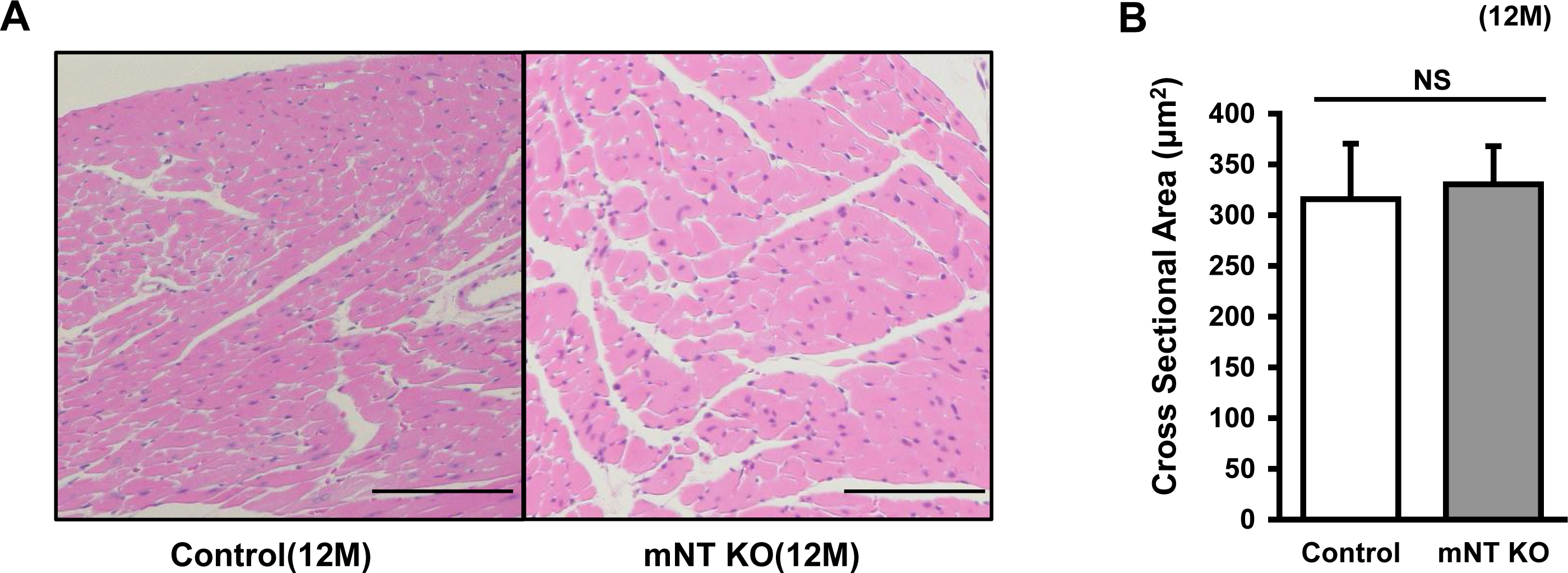
Representative Histological Images of the Heart. (A) Representative Hematoxylin and eosin (HE) stains from 12-month old control mice and 12-month old mitoNEET-knockout mice. Scale Bar, 100μm. (B) Summary data for cross sectional area. n=3 for each.

**Appendix Figure S6.**
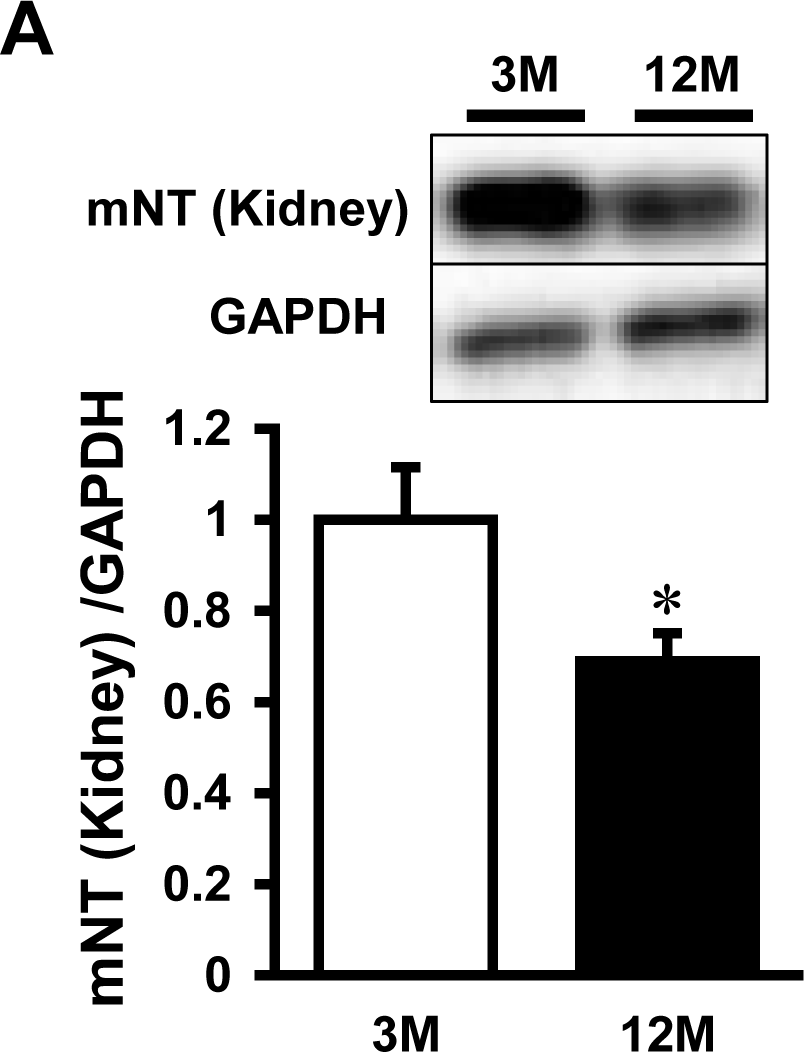
Expression of mitoNEET in the kidney of 12-Month Old C57B6/J Mice. (A) Representative immunoblot and summary data of mitoNEET protein expression normalized to GAPDH in the kidneys of 3 and 12-month old mice. Data are shown as the mean ± SE. n=5-6. *P<0.05 vs. 3M. M, month; mNT, mitoNEET; GAPDH, glyceraldehyde phosphate dehydrogenase.

**Appendix Table S1.**

| Identified Protein | Score | Molecular weight (kDa) | Expected value | NCBI accession number |
| --- | --- | --- | --- | --- |
| <band around 100 kDa> |  |  |  |  |
| Transferrin receptor protein 1 | 131 | 85274 | 9.2e-009 | cd09848 |

MASCOT Scores and National Center for Biotechnology Information (NCBI) Accession Numbers of Proteins. Generated by the MASCOT database (Matrix Science, Boston, MA).

**Appendix Table S2.**
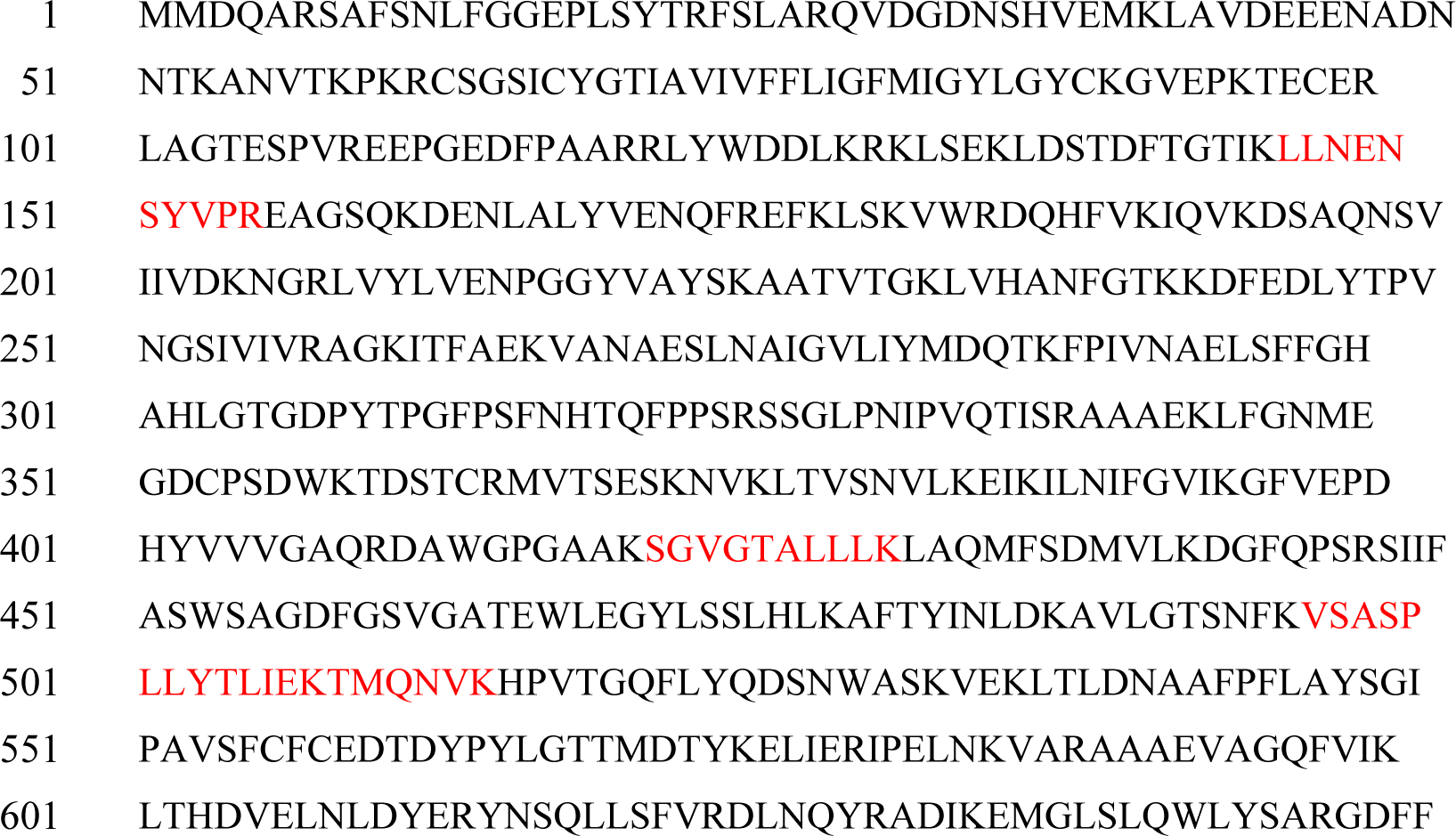
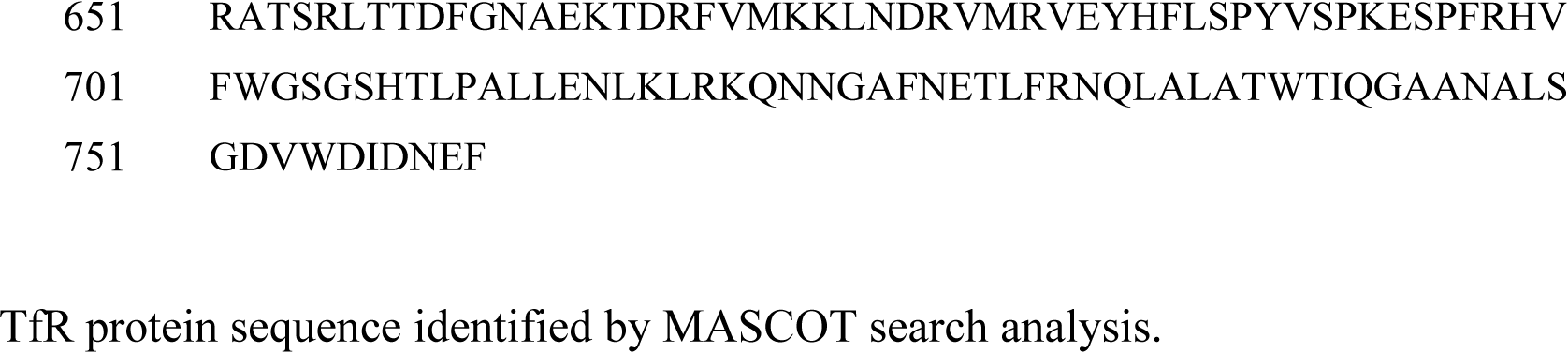
Protein sequence coverage: 5% Matched peptides shown in **red**.

## Reference

Anderson CP, Shen M, Eisenstein RS, Leibold EA (2012) Mammalian iron metabolism and its control by iron regulatory proteins. Biochim Biophys Acta 1823: 1468–83

Anderson GJ, Vulpe CD (2009) Mammalian iron transport. Cell Mol Life Sci 66: 3241– 61

Arasaki K, Shimizu H, Mogari H, Nishida N, Hirota N, Furuno A, Kudo Y, Baba M, Baba N, Cheng J, Fujimoto T, Ishihara N, Ortiz-Sandoval C, Barlow LD, Raturi A, Dohmae N, Wakana Y, Inoue H, Tani K, Dacks JB et al. (2015) A role for the ancient SNARE syntaxin 17 in regulating mitochondrial division. Dev Cell 32: 304–17

Bagh MB, Thakurta IG, Biswas M, Behera P, Chakrabarti S (2011) Age-related oxidative decline of mitochondrial functions in rat brain is prevented by long term oral antioxidant supplementation. Biogerontology 12: 119–31

Branda SS, Cavadini P, Adamec J, Kalousek F, Taroni F, Isaya G (1999) Yeast and human frataxin are processed to mature form in two sequential steps by the mitochondrial processing peptidase. J Biol Chem 274: 22763–9

Christiansen LB, Dela F, Koch J, Hansen CN, Leifsson PS, Yokota T (2015) Impaired cardiac mitochondrial oxidative phosphorylation and enhanced mitochondrial oxidative stress in feline hypertrophic cardiomyopathy. Am J Physiol Heart Circ Physiol 308: H1237–47

Dai DF, Rabinovitch PS (2009) Cardiac aging in mice and humans: the role of mitochondrial oxidative stress. Trends Cardiovasc Med 19: 213–20

Das A, Nag S, Mason AB, Barroso MM (2016) Endosome-mitochondria interactions are modulated by iron release from transferrin. J Cell Biol 214: 831–45

Dranka BP, Hill BG, Darley-Usmar VM (2010) Mitochondrial reserve capacity in endothelial cells: The impact of nitric oxide and reactive oxygen species. Free Radic Biol Med 48: 905–14

Fukushima A, Kinugawa S, Takada S, Matsushima S, Sobirin MA, Ono T, Takahashi M, Suga T, Homma T, Masaki Y, Furihata T, Kadoguchi T, Yokota T, Okita K, Tsutsui H (2014) (Pro)renin receptor in skeletal muscle is involved in the development of insulin resistance associated with postinfarct heart failure in mice. Am J Physiol Endocrinol Metab 307: E503–14

Ganz T (2013) Systemic iron homeostasis. Physiol Rev 93: 1721–41

Gkouvatsos K, Papanikolaou G, Pantopoulos K (2012) Regulation of iron transport and the role of transferrin. Biochim Biophys Acta 1820: 188–202

Hey-Mogensen M, Hojlund K, Vind BF, Wang L, Dela F, Beck-Nielsen H, Fernstrom M, Sahlin K (2010) Effect of physical training on mitochondrial respiration and reactive oxygen species release in skeletal muscle in patients with obesity and type 2 diabetes. Diabetologia 53: 1976–85

Ichikawa Y, Bayeva M, Ghanefar M, Potini V, Sun L, Mutharasan RK, Wu R, Khechaduri A, Jairaj Naik T, Ardehali H (2012) Disruption of ATP-binding cassette B8 in mice leads to cardiomyopathy through a decrease in mitochondrial iron export. Proc Natl Acad Sci U S A 109: 4152–7

Ichikawa Y, Ghanefar M, Bayeva M, Wu R, Khechaduri A, Naga Prasad SV, Mutharasan RK, Naik TJ, Ardehali H (2014) Cardiotoxicity of doxorubicin is mediated through mitochondrial iron accumulation. J Clin Invest 124: 617–30

Ide T, Tsutsui H, Kinugawa S, Utsumi H, Takeshita A (1999) Amiodarone protects cardiac myocytes against oxidative injury by its free radical scavenging action. Circulation 100: 690–2

Ikeuchi M, Matsusaka H, Kang D, Matsushima S, Ide T, Kubota T, Fujiwara T, Hamasaki N, Takeshita A, Sunagawa K, Tsutsui H (2005) Overexpression of mitochondrial transcription factor a ameliorates mitochondrial deficiencies and cardiac failure after myocardial infarction. Circulation 112: 683–90

Isaya G (2014) Mitochondrial iron-sulfur cluster dysfunction in neurodegenerative disease. Front Pharmacol 5: 29

Khechaduri A, Bayeva M, Chang HC, Ardehali H (2013) Heme levels are increased in human failing hearts. J Am Coll Cardiol 61: 1884–93

Kinugawa S, Tsutsui H, Hayashidani S, Ide T, Suematsu N, Satoh S, Utsumi H, Takeshita A (2000) Treatment with dimethylthiourea prevents left ventricular remodeling and failure after experimental myocardial infarction in mice: role of oxidative stress. Circ Res 87: 392–8

Kusminski CM, Holland WL, Sun K, Park J, Spurgin SB, Lin Y, Askew GR, Simcox JA, McClain DA, Li C, Scherer PE (2012) MitoNEET-driven alterations in adipocyte mitochondrial activity reveal a crucial adaptive process that preserves insulin sensitivity in obesity. Nat Med 18: 1539–49

Kwong LK, Sohal RS (2000) Age-related changes in activities of mitochondrial electron transport complexes in various tissues of the mouse. Arch Biochem Biophys 373: 16–22

Li H, Kumar Sharma L, Li Y, Hu P, Idowu A, Liu D, Lu J, Bai Y (2013) Comparative bioenergetic study of neuronal and muscle mitochondria during aging. Free Radic Biol Med 63: 30–40

Matsushima S, Ide T, Yamato M, Matsusaka H, Hattori F, Ikeuchi M, Kubota T, Sunagawa K, Hasegawa Y, Kurihara T, Oikawa S, Kinugawa S, Tsutsui H (2006) Overexpression of mitochondrial peroxiredoxin-3 prevents left ventricular remodeling and failure after myocardial infarction in mice. Circulation 113: 1779–86

Minotti G, Menna P, Salvatorelli E, Cairo G, Gianni L (2004) Anthracyclines: molecular advances and pharmacologic developments in antitumor activity and cardiotoxicity. Pharmacol Rev 56: 185–229

Oliveira F, Rocha S, Fernandes R (2014) Iron metabolism: from health to disease. J Clin Lab Anal 28: 210–8

Paddock ML, Wiley SE, Axelrod HL, Cohen AE, Roy M, Abresch EC, Capraro D, Murphy AN, Nechushtai R, Dixon JE, Jennings PA (2007) MitoNEET is a uniquely folded 2Fe 2S outer mitochondrial membrane protein stabilized by pioglitazone. Proc Natl Acad Sci U S A 104: 14342–7

Paradkar PN, Zumbrennen KB, Paw BH, Ward DM, Kaplan J (2009) Regulation of mitochondrial iron import through differential turnover of mitoferrin 1 and mitoferrin 2. Mol Cell Biol 29: 1007–16

Perdomini M, Belbellaa B, Monassier L, Reutenauer L, Messaddeq N, Cartier N, Crystal RG, Aubourg P, Puccio H (2014) Prevention and reversal of severe mitochondrial cardiomyopathy by gene therapy in a mouse model of Friedreich’s ataxia. Nat Med 20: 542–7

Richardson DR, Lane DJ, Becker EM, Huang ML, Whitnall M, Suryo Rahmanto Y, Sheftel AD, Ponka P (2010) Mitochondrial iron trafficking and the integration of iron metabolism between the mitochondrion and cytosol. Proc Natl Acad Sci U S A 107: 10775–82

Rosca MG, Hoppel CL (2010) Mitochondria in heart failure. Cardiovasc Res 88: 40–50

Srinivasan V, Pierik AJ, Lill R (2014) Crystal structures of nucleotide-free and glutathione-bound mitochondrial ABC transporter Atm1. Science 343: 1137–40

Suematsu N, Tsutsui H, Wen J, Kang D, Ikeuchi M, Ide T, Hayashidani S, Shiomi T, Kubota T, Hamasaki N, Takeshita A (2003) Oxidative stress mediates tumor necrosis factor-alpha-induced mitochondrial DNA damage and dysfunction in cardiac myocytes. Circulation 107: 1418–23

Sugiyama S, Takasawa M, Hayakawa M, Ozawa T (1993) Changes in skeletal muscle, heart and liver mitochondrial electron transport activities in rats and dogs of various ages. Biochem Mol Biol Int 30: 937–44

Takada S, Masaki Y, Kinugawa S, Matsumoto J, Furihata T, Mizushima W, Kadoguchi T, Fukushima A, Homma T, Takahashi M, Harashima S, Matsushima S, Yokota T, Tanaka S, Okita K, Tsutsui H (2016) Dipeptidyl peptidase-4 inhibitor improved exercise capacity and mitochondrial biogenesis in mice with heart failure via activation of glucagon-like peptide-1 receptor signalling. Cardiovasc Res 111: 338–47

Tonkonogi M, Sahlin K (1997) Rate of oxidative phosphorylation in isolated mitochondria from human skeletal muscle: effect of training status. Acta Physiol Scand 161: 345–53

Vaubel RA, Isaya G (2013) Iron-sulfur cluster synthesis, iron homeostasis and oxidative stress in Friedreich ataxia. Mol Cell Neurosci 55: 50–61

Vigani G, Tarantino D, Murgia I (2013) Mitochondrial ferritin is a functional iron-storage protein in cucumber (Cucumis sativus) roots. Front Plant Sci 4: 316

Wiley SE, Murphy AN, Ross SA, van der Geer P, Dixon JE (2007a) MitoNEET is an iron-containing outer mitochondrial membrane protein that regulates oxidative capacity. Proc Natl Acad Sci U S A 104: 5318–23

Wiley SE, Paddock ML, Abresch EC, Gross L, van der Geer P, Nechushtai R, Murphy AN, Jennings PA, Dixon JE (2007b) The outer mitochondrial membrane protein mitoNEET contains a novel redox-active 2Fe-2S cluster. J Biol Chem 282: 23745–9

Wittig I, Braun HP, Schagger H (2006) Blue native PAGE. Nat Protoc 1: 418–28

Yamamoto A, Takeya R, Matsumoto M, Nakayama KI, Sumimoto H (2013) Phosphorylation of Noxo1 at threonine 341 regulates its interaction with Noxa1 and the superoxide-producing activity of Nox1. FEBS J 280: 5145–59

